# SUMO monoclonal antibodies vary in sensitivity, specificity, and ability to detect SUMO conjugate types

**DOI:** 10.1101/2022.03.19.484974

**Authors:** Alexander J. Garvin, Alexander J. Lanz, Joanna R. Morris

**Affiliations:** Birmingham Centre for Genome Biology and Institute of Cancer and Genomic Sciences, College of Medical and Dental Schools, University of Birmingham, B15 2TT, UK

## Abstract

Monoclonal antibodies (MAb) to members of the Small Ubiquitin-like modifier (SUMO) family are essential tools in the study of cellular SUMOylation. However, many reagents are poorly validated, and the choice of which antibody to use for which detection format is without an evidence base. Here we test twenty-four anti-SUMO monoclonal antibodies for sensitivity and specificity to SUMO1-4 in monomeric and polymeric states in dot-blots, immunoblots, immunofluorescence and immunoprecipitation. We find substantial variability between SUMO MAbs for different conjugation states, detection of increased SUMOylation in response to thirteen different stress agents and as enrichment reagents for analysis of SUMOylated RanGAP1 or KAP1. All four anti-SUMO4 monoclonal antibodies tested cross-reacted with SUMO2/3, and several SUMO2/3 monoclonal antibodies cross-reacted with SUMO4. ~10% of tested monoclonal antibodies produced specific results across multiple applications.

## Introduction

The SUMO family is made up of three conjugated members (SUMO1-3), a non-conjugatable SUMO4 (Varejao et al., 2020) and SUMO5/SUMO1P1, which has restricted tissue expression (Liang et al., 2016). SUMO1-3 are processed into mature conjugatable forms through the removal of the extreme C-terminal residues (Mikolajczyk et al., 2007). SUMO1 and SUMO2/3 form two distinct branches of the SUMO system, which use the same conjugation machinery (Wang et al., 2014, Bouchard et al., 2021). SUMO isoforms can be conjugated as monomers, multi-monomers and polymers and can form multiple internal lysine linkage types, including branching and mixed chains comprised of different SUMO isoforms and other Ub/Ubls (Morris and Garvin, 2017). Conjugation of SUMO (SUMOylation) is essential for several cellular processes, including transcription, DNA replication, mitosis, genome stability and immunity (Ryu and Hochstrasser, 2021)(Chang et al., 2021, Boulanger et al., 2021, Kroonen and Vertegaal, 2021, Abbas, 2021)(Garvin and Morris, 2017). Transient up-regulation of SUMOylation is associated with responses to cellular stress (Ilic et al., 2022). The modification can alter protein localisation, activity, turnover, and protein interactions (Lascorz et al., 2021, Pichler et al., 2017). SUMOylation is a transient process often confined to a subset of the target protein that may be spatially and temporally restricted. SUMO proteases (SENP1-7) mediate deconjugation. The balance of SUMOylation/deSUMOylation is essential for many cellular processes (Kunz et al., 2018, Garvin, 2019, Tokarz and Wozniak, 2021).

Advances in proteomic analysis of SUMO conjugation have enhanced cataloguing of the global SUMOylome (Hendriks et al., 2017, Tammsalu et al., 2014, Lamoliatte et al., 2017) with further adaptions reducing dependency on over-expressed epitope-tagged SUMO (Becker et al., 2013, Hendriks et al., 2018). Enrichment with SUMO interacting trap proteins (Bruderer et al., 2011, Da Silva-Ferrada et al., 2013) or BioID (Pirone et al., 2017) have added further to our understanding of the SUMOylated proteome. Detection of SUMO conjugation is challenging due to the small proportion of modified substrate, its transient, often context-dependent nature and the rapid deconjugation by SENP enzymes. SUMOylation studies rely on the detection of endogenous SUMO family members using antibodies.

Approximately one hundred SUMO1-4 monoclonal antibodies are commercially available (Supplemental Table 1). Of these, a minority are cited (Supplemental Figure 1a), and most are poorly or not validated (Supplemental Figure 1b-c). An ongoing expansion in the number of available antibodies, coupled with poor or no validation has the potential to contribute significant confusion and difficulty in reproducibility in the biomedical sciences. Antibodies are often highlighted as a major contributor to the reproducibility crisis in research (Goodman, 2018, Baker, 2015).

A previous methodical attempt to re-validate seven reported neuronal SUMO1 conjugated proteins using a HA-SUMO1 knock-in mouse failed to confirm SUMO1 conjugation for any of these substrates (Tirard et al., 2012, Daniel et al., 2017). While differences in methodology, expression levels and animal models may explain some of these issues, significant deficiencies in available SUMO1 antibodies were noted (Wilkinson et al., 2017, Daniel et al., 2018). Additionally, anecdotal experience of variability in anti-SUMO antibody reagents to detect SUMO conjugation following ionising radiation treatment (Garvin et al., 2019b) leads us here to catalogue the specificity and sensitivity of some the most commonly used SUMO MAbs. Here we provide a comprehensive analysis of twenty-four SUMO monoclonal antibodies and highlight the strengths and weaknesses of these reagents.

## Results

We chose 24 MAbs from the 93 SUMO1-4 MAbs commercially available at the time of writing (Supplemental Table 1); 9 raised against SUMO1, 11 against SUMO2/3 and 4 against SUMO4. They were selected based on high citations from the CiteAb database as a proxy for research community usage, and each had validation data available on the manufacturer’s websites (Helsby et al., 2014). Antibodies were raised in mice, rabbits and rats and used a variety of immunogens, including recombinant GST-SUMO, untagged SUMO, and peptides. With three exceptions (SUMO2+3 8A2, SUMO2+3 12F3, SUMO1 21C7) the antibody epitopes had not been mapped or the identity of the peptide immunogen was proprietary (Supplemental Table 1).

In this study, antibodies were tested at 1 μg/mL except for recombinant antibodies (EPR300, EPR4602, EPR7163, JJ-085 and ARC1382) or antibodies from Cell Signalling Technologies (C9H1 and 18H8), which are supplied at lot-specific dilutions. In these exceptions, antibodies were diluted at 1:1000 in 5% milk. As some of the MAbs used are available from multiple vendors, they are referred to by clone name rather than catalogue number throughout.

### Sensitivities and specificity of MAbs for monomeric SUMO isoforms

To test the sensitivity and specificity of the antibodies, we generated recombinant SUMO1-4 purified from *E. coli* (rSUMO). As most of the antibodies raised against SUMO proteins are generated against the immature (Pro)SUMO isoforms (Supplemental Table 1), we assayed both immature and mature (terminating in GG) forms. Unlike SUMO1-3, SUMO4 is not processed into a mature form (Owerbach et al., 2005). For SUMO4, we purified both WT SUMO4 and the M55V variant (rs237025). The polymorphism underlying SUMO4 M55V is prevalent and associated with several human pathologies, including diabetes (Guo et al., 2004).

Each SUMO1 antibody was tested by dot blot of serially diluted rSUMO1. As SUMO1 shares just 50% amino acid identity with SUMO2-4, we did not test cross-reactivity with SUMO2/3 isoforms (Figure 1a). A range of sensitivities was clear. 4D12 showed a weak signal, for example, while Y299 was saturating (Figure 1b). We detected no differences in the ability of the MAbs to detect pro-SUMO1 *Vs* mature forms of rSUMO1 (Figure 1b). Antibodies raised against SUMO2/3 showed a range of sensitivities to recombinant SUMO2 and SUMO3, from almost no signal (EPR4602, EPR300, SM23/496) to a strong signal even at low concentrations and short exposures (8A2 and 12F3) with no differences in the ability to detect pro and mature SUMO2/3 forms (Figure 1c). None showed a preference for SUMO2 or SUMO3. The antibody 2H8, and to a lesser degree 852908, detected rSUMO4 by dot blot. The four tested antibodies raised to SUMO4 detected rSUMO4 to varying degrees, with none showing differential detection of the M55V variant. All showed cross-reaction to rSUMO2/3 (Figure 1c).

**Figure 1.**
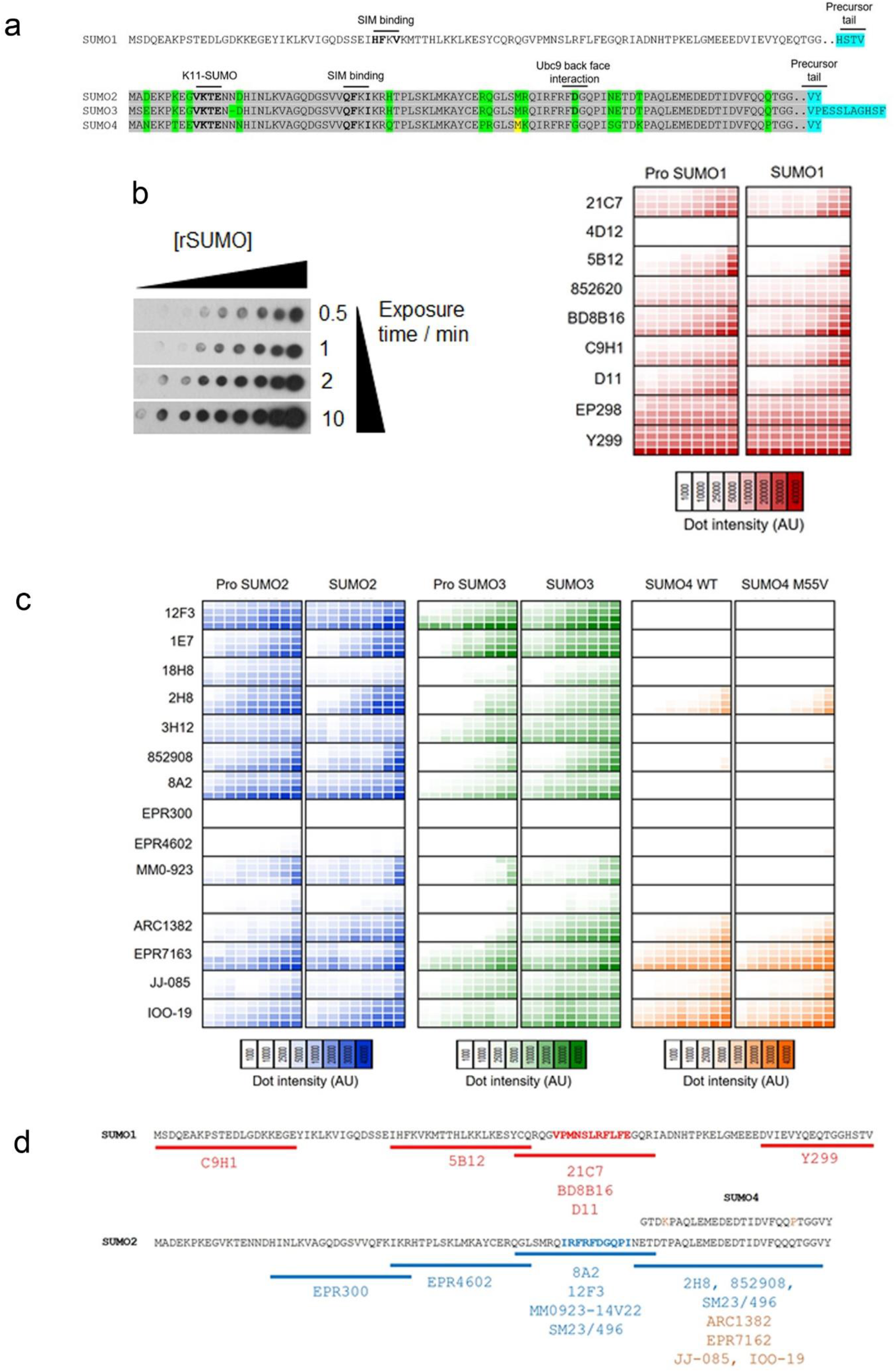
**a.** Illustration of SUMO1-4 sequence with known epitopes. Amino acid sequences of human SUMO1, SUMO2, SUMO3 and SUMO4. SIM (SUMO Interacting Motif) contacting residues, Ubc9 back face interacting residues and precursor tail residues are indicated. Conserved residues between SUMO2-4 are highlighted in grey while non conserved residues are highlighted in green. The variable M55 residue in SUMO4 is highlighted in yellow. The precursor tail residues removed in the mature SUMO proteins are highlighted in blue. **b.** Heat map of SUMO1 MAbs detecting rSUMO1. Heat map of dot blot signal for SUMO1 MAbs. Signal intensity (by chemiluminescent detection on X-ray film) is shown by heat map, with deeper colour signifying more intense chemiluminescent signal, for four exposure time points (0.5, 1, 2 and 10 minutes) as an average of three independent repeats. A representative dot blot is shown with increasing concentration of rSUMO on the horizontal and increasing film exposure time on the vertical. **c.** Heat map of SUMO2/3/4 MAbs detecting rSUMO/3/4. As for 1B, but each MAb tested against SUMO2 and SUMO3 (Pro and mature forms) and SUMO4 (WT and M55V variant). **d.** Epitope mapping of SUMO MAbs. Partially overlapping SUMO1, SUMO2 or SUMO4 peptides used for competition assays with loss of detection (by dot blot or immunoblot of SUMO conjugates) indicating location of individual MAb epitopes. SUMO1 region highlighted in red shows the mapped 21C7 epitope and SUMO2 region highlighted in blue shows the 8A2 epitope. MAbs not shown could not be mapped.

To better understand antibody cross-reactivity, we used peptide competition assays and were able to provide approximate epitope locations for 18 out of 24 MAbs (Figure 1d). The mapping also confirmed the location of 21C7 and 8A2 epitopes which have previously been mapped (Becker et al., 2013, Subramonian et al., 2014). All four antibodies raised to SUMO4 detected the C terminus of SUMO4, a region of high (92%) conservation with SUMO2/3 (Figure 1a), and the two most SUMO4 cross-reactive SUMO2/3 clones (2H8 and 852908) also recognised this region.

Overall, we find a wide range of sensitivities for monomeric SUMO isoforms, little specificity for pro *Vs* mature SUMO isoforms, and antibodies raised to SUMO4 cross react with SUMO2/3.

### Varied sensitivity and specificity for SUMO isoforms in cell lysates

Next, we sought to determine specificity and selectivity against cellular SUMO conjugates. We generated cells able to induce cDNAs for SUMO1, SUMO3 and SUMO4 containing synonymous mutations rendering them insensitive to siRNA depleting the endogenous SUMO transcript, while endogenous SUMO2 was targeted using siRNA to the 3’ UTR (Supplemental Figure 2a). Simultaneous siRNA depletion and Dox addition allowed a degree of replacement of endogenous SUMO with FLAG-tagged variants that could be probed with the well-established anti-FLAG antibody, M2, as a comparison.

Using this approach, we found antibodies raised to SUMO1: 21C7, BD8B16 and Y299 were sensitive and specific; the remaining had less sensitivity (Figure 2a & b). D11 and EP298 showed some reduction on siRNA treatment, but also detected non-specific bands. Fractionation of cell lysates confirmed these bands to be predominantly cytoplasmic and not detected in chromatin where most SUMO1 conjugates are found (Supplemental Figure 2b). 852620 detected proteins that were unchanged by prior SUMO1 siRNA treatment. The banding pattern of this clone was also unchanged when cells were treated with SUMO1 E1 inhibitor ML-792 (He et al., 2017), the NEDD8 E1 inhibitor MLN4924 and were not increased by the ubiquitin E1 inhibitor TAK-243, which causes increased SUMOylation (Sha et al., 2019) (Supplemental Figure 2c).

**Figure 2.**
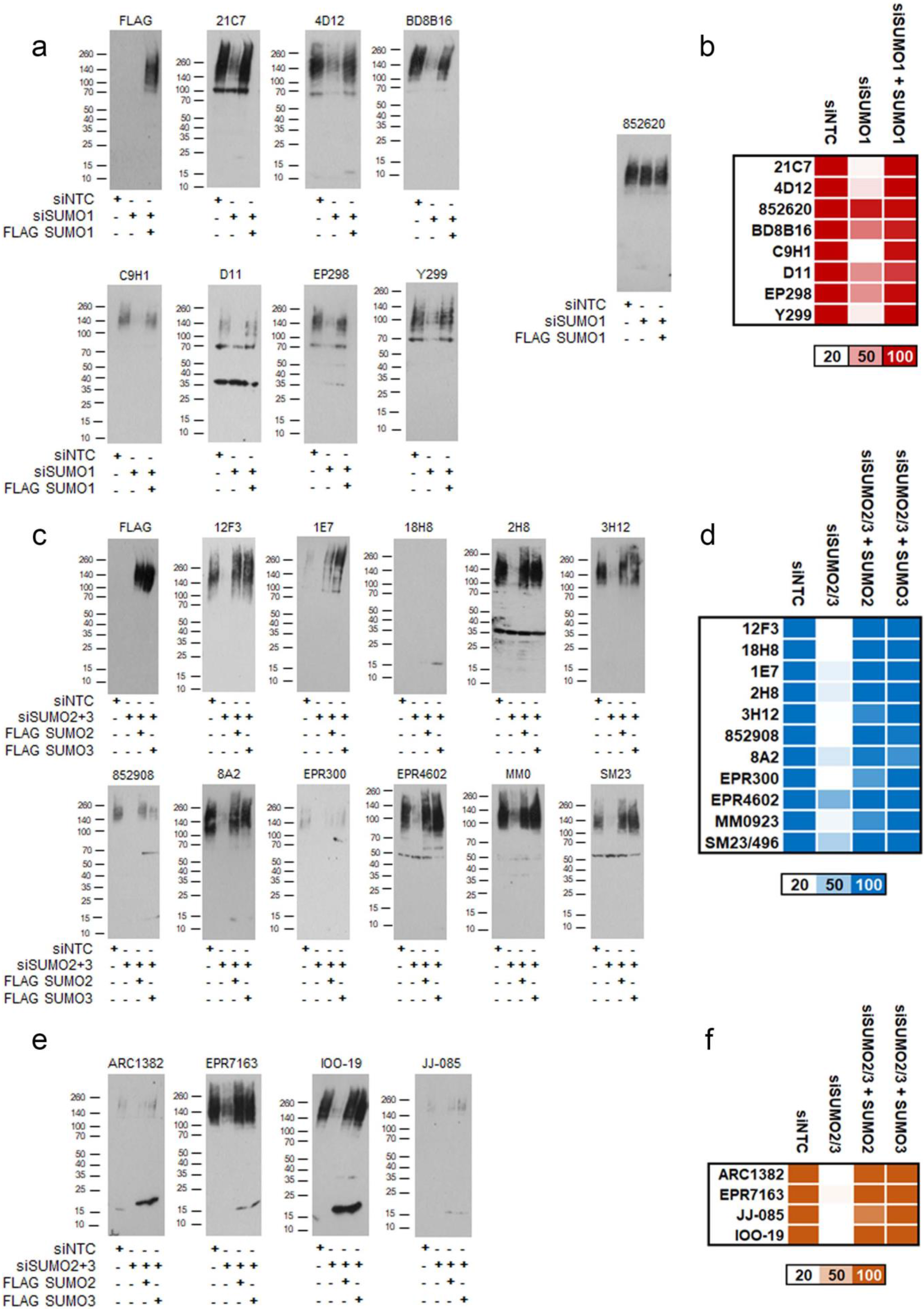
**a.** U2OS FLAG-SUMO1 treated with siNTC, siSUMO1 or siSUMO1 with doxycycline for 48 hours in 6 well plates were lysed in 400 μL 6x Laemmli buffer. 20 μL of lysate per lane were separated by 4-20% SDS-PAGE and immunoblotted with indicated MAbs. All blots are shown at 1 min exposure. **b.** Heat map of SUMO1 conjugate intensity (total conjugates including free SUMO) shown as % of siNTC at 1 minute exposure. **c.** As for 2a but using FLAG-SUMO2 and FLAG-SUMO3 expressing U2OS cells. Cells were treated with siRNA to both SUMO2 and SUMO3. **d.** Heat map of SUMO2/3 as for 2b. **e.** Lysates from 2c probed with SUMO4 MAbs. **f.** As for 2b but with SUMO4 MAbs.

Antibodies raised to SUMO2/3 and SUMO4 showed reduced signal when probing lysates from SUMO2/3 siRNA depleted cells and showed similar detection of FLAG-SUMO2 and FLAG-SUMO3 (Figure 2c & d). Antibodies raised to SUMO4 detected high molecular weight FLAG-SUMO2 and SUMO3 (Figure 2e & f) and strongly detected the HA-FLAG-SUMO4. Similarly, anti-SUMO2/3 antibodies 2H8 and 852908 showed strong detection of HA-FLAG-SUMO4. In contrast to the dot blot data, we also detected varying degrees of HA-FLAG-SUMO4 cross-reaction by several SUMO2/3 MAbs (Supplemental Figure 2d-g). Thus, antibodies raised to SUMO4 detect SUMO2/3, and some raised to SUMO2/3 detect SUMO4.

### Differences in the ability to detect free and conjugated SUMO

We quantified the % free SUMO isoforms (~18 KDa band) in lysates from A427, CAL51, CALU6, HCT116, HEK293 and U2OS cells detected by each antibody. The majority did not bind to free SUMO in western blots, while for a minority, 10-20% of the total SUMO signal could be accounted for by this band (Figure 3a & b and Supplemental Figure 3a). Next, we tested detection of recombinant monomeric, dimeric and polySUMO2 by antibodies raised to SUMO2/3/4 (Figure 3a & b and Supplemental Figure 3b. Some exhibited a preference for monomeric SUMO2 over polymeric forms (12F3, 2H8, 852908, 8A2 and SM23/496), others showed less preference (1E7, 3H12, EPR4602, MM093-14V22). Some bound poorly to high molecular weight SUMO2 conjugates (18H8 and EPR300). All four antibodies raised to SUMO4 bound to recombinant SUMO2 monomers, dimers, and polymers (Figure 3b & c).

**Figure 3.**
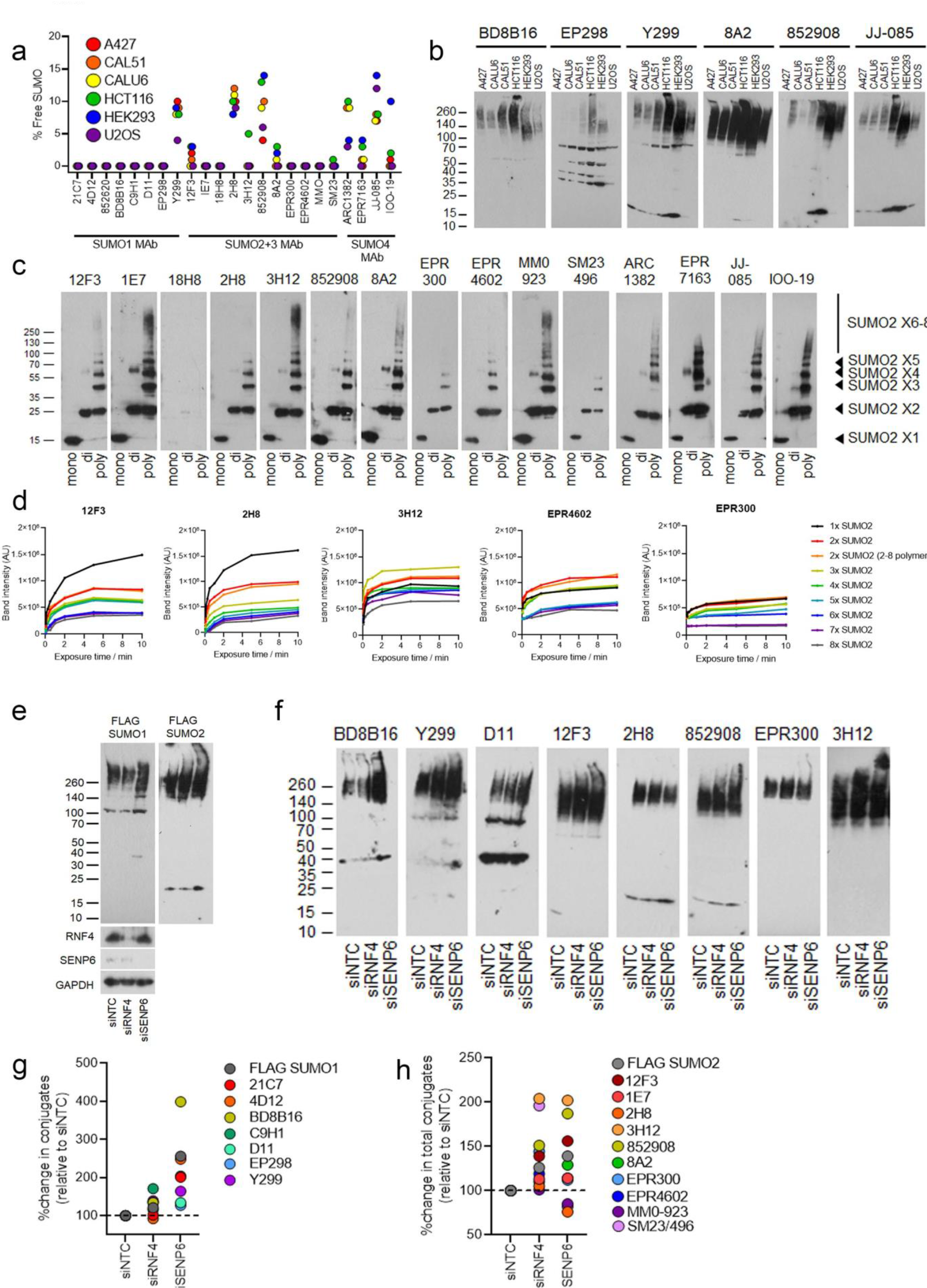
**a.** The % free SUMO relative to the total SUMO signal is shown for each of six cell lines, band intensity was calculated from multiple exposures. **b.** Representative blots of lysates from indicated cell lines, the band at ~15 KDa is free SUMO. Images for all cell lines and exposures can be found in Supplemental figure 3A. **c.** Immunoblots of SUMO2 monomers, dimers, and polymers (2-8x SUMO2) at 200 ng / lane resolved on 4-20% SDS PAGE gels. Images are from various times of film exposures. All exposures times for each MAb can be found in in Supplemental figure 3B. **d.** Quantification of representative experiments from 3C from multiple exposure times. **e.** Immunoblots of U2OS FLAG-SUMO1 or FLAG-SUMO2 cells siRNA depleted and dox treated for 48 hours to replace endogenous SUMO with FLAG-SUMO. Depletion efficiencies of RNF4 and SENP6 are shown. **f.** Representative immunoblots FLAG-SUMO complemented U2OS lysates from 3E. **g.** The % change in total SUMO1 conjugates following siRNA depletion of RNF4 and SENP6 relative to non-targeting (NTC) siRNA control. **h.** As for 3g but with SUMO2/3 Mabs.

To further explore potential variability for the analysis of cellular conjugates, we increased SUMO polymer levels by depleting the polySUMO targeted ubiquitin ligase RNF4 and polySUMO specific protease SENP6 in FLAG-SUMO1 or FLAG-SUMO2 complemented cells. The anti-FLAG signal showed the expected shifts in molecular weight associated with highly modified SUMO conjugates following these depletions, as did clones BD8B16, C9H1 and 3H12. Some antibodies, D11, EP298, 2H8 and MM0923, showed almost no change in conjugate levels (Figure 3d & e). Collectively this data shows antibodies vary in their ability to distinguish different forms of SUMO.

### Detection of SUMO conjugates on stress

Global proteomic evidence suggests that different stresses result in both alterations in substrates and types of SUMO modifications (Hendriks et al., 2015, Hendriks et al., 2018). We tested thirteen different agents that included genotoxic (IR and CPT), replicative (HU), oxidative (H2O2), proteotoxic (MG132, CB-5083 and 17-AAG), osmotic (Sorbitol), thermal (Heat Shock) and mitotic (ICRF-193) stresses. The conditions tested resulted in varying degrees of FLAG-SUMO conjugate changes. Treatment with the proteasome inhibitor, MG132, resulted in increasing conjugates of SUMO1 and SUMO2 using most tested antibodies (Figure 4a-d and Supplemental Figure 4a & b). Three antibodies detected large increases in SUMO conjugates following irradiation (4D12, 1E7 and 3H12), others did not, or detected a decrease in conjugates (2H8, MM0-923 and SM23/496). The conjugate pattern of the SUMO4 MAbs resembled the pattern detected by clones raised to SUMO2/3 rather than HA-FLAG SUMO4 detected by M2 (Figure 4e & f and Supplemental Figure 4c).

**Figure 4.**
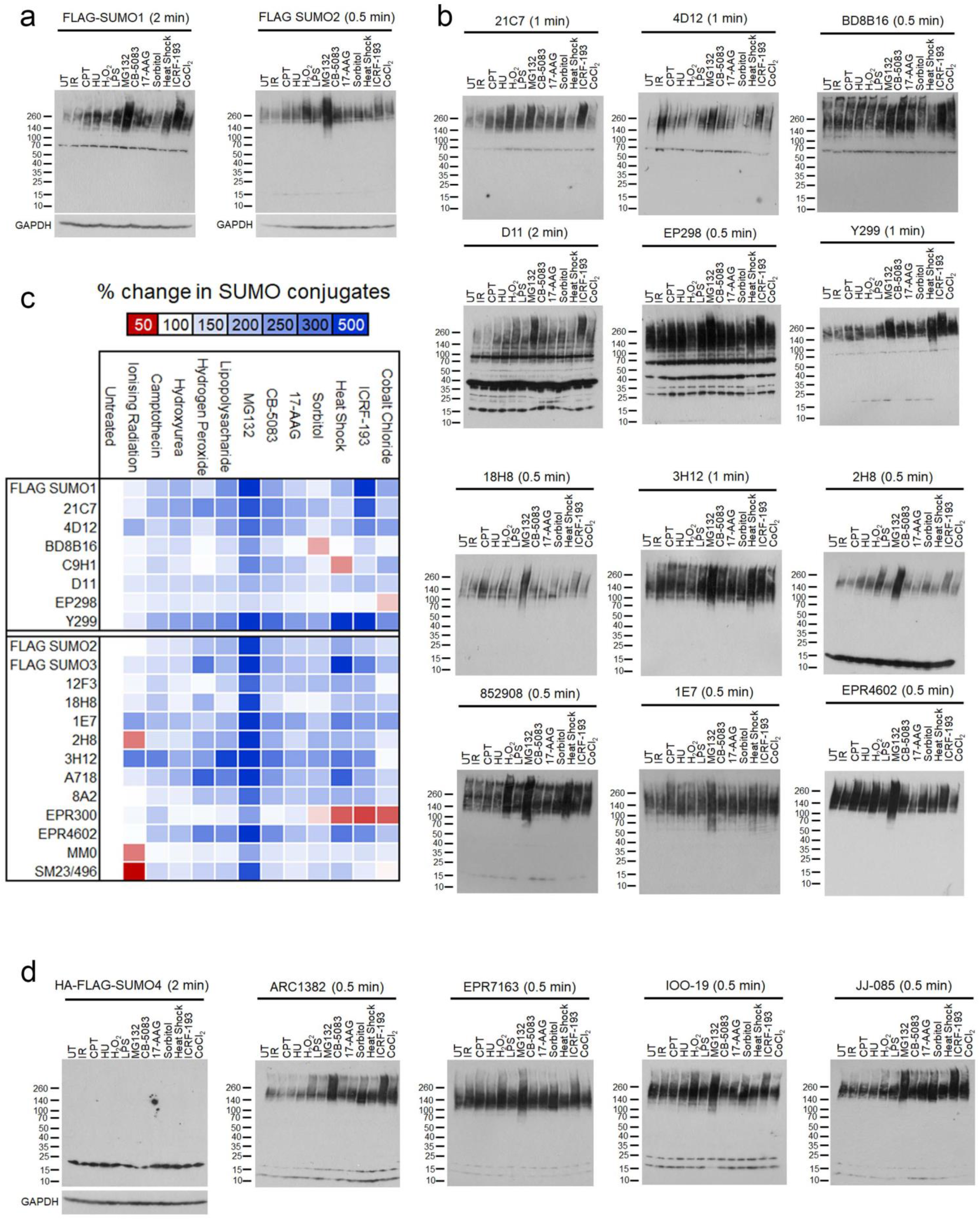
**a.** U2OS plated in 6 well plates, siRNA depleted of endogenous SUMO1 or SUMO2/3 and doxycline treated to induce FLAG-SUMO1 or FLAG-SUMO2 for 48 hours prior to treatment with indicated stress agents. UT (Untreated control), IR (Ionising Radiation, 4 Gy, 1 hr post), CPT (Camptothecin 1 μM, 3 hr), HU (Hydroxyurea 5 mM, 3 hr), H2O2 (Hydrogen Peroxide 125 μM, 15 minutes), LPS (Lipopolysaccharide 1000 ng, 3 hr), MG132 (5 μg/mL, 3 hr), CB-5083 (0.1 μM, 3 hr), 17-AAG (5 μM, 3 hr), Sorbitol (500 mM, 30 min), heat shock (42 °C, 10 min), ICRF-193 (100 μM, 1 hr). Lysates were made by direct lysis in 6x Laemlii buffer, 20 μL of a total of 400 μL per 6 well was run per lane. Exposure time is shown, all exposures are shown in Supplemental Figure 4a and 4b. **b.** Representative immunoblots of lysates from 4a with SUMO1 (21C7, 4D12, BD8B16, D11, EP298 and Y299) and SUMO2/3 (18H8, 3H12, 2H8, 852908, 1E7 and EPR4602) with exposure times shown. All exposures and MAbs are shown in Supplemental Figure 4a and 4b. **c.** Heat map summarising changes in SUMO conjugates after stress for each SUMO1 or SUMO2/3 MAb. % Change in SUMO conjugates are calculated from three experimental repeats per MAb using densitometry of SUMO conjugates migrating above 70 kDa (free SUMO not included) as a % change relative to the untreated control lane for each immunoblot. The mean for each condition and MAb is shown. **d.** U2OS expressing HA-FLAG-SUMO4 treated as for 4a. All exposures are shown in Supplemental Figure 2b.

### Specificity of MAbs for indirect immunofluorescence

We next tested the ability of the clones to detect SUMO subcellular localisation by immunofluorescence in fixed U2OS cells. In siRNA complemented U2OS, exogenous FLAG-SUMO1 and FLAG-SUMO2 predominantly localised to the nucleus and HA-FLAG-SUMO4 localised to both the nucleus and cytoplasm (Figure 5a). In contrast, the antibodies detected variable patterns, with some showing a diffuse nuclear signal and others a concentration of dot-like structures, which co-localised with PML (Figure 5b & c). To confirm the specificity, we also stained siRNA depleted cells. Staining patterns were reduced by siRNA depletion for some clones (e.g. SUMO1: BD8B16, Y299 SUMO2/3 SUMO2/3 3H1, 8A2) others were not (e.g., SUMO2/3 2Hb and EPR300) (Figure 5e & f). The specificity of staining was also affected by the fixation and extraction conditions used (Supplemental Figure 5a). SUMO4 MAbs exhibited predominantly nuclear signals and depletion of SUMO2 and SUMO3, but not SUMO4, decreased fluorescence signals (Figure 5d and g). Taken together, these data show that the antibodies raised to SUMO isoforms vary greatly in their sensitivity and specificity for SUMO isoforms and their conjugates by indirect immunofluorescence.

**Figure 5.**
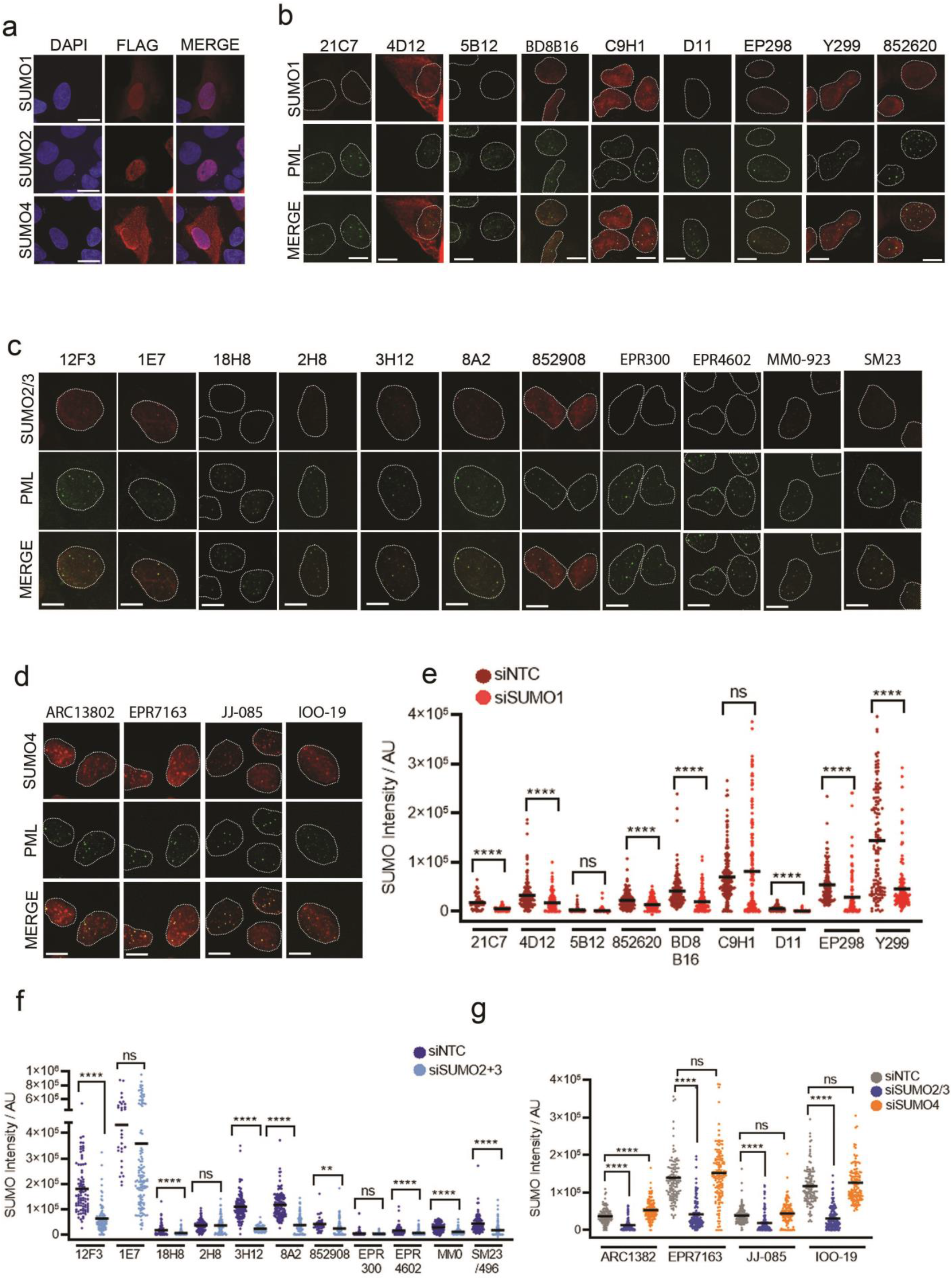
**a.** Localisation of FLAG-SUMO1, FLAG-SUMO2 and HA-FLAG SUMO4 in U2OS by indirect immunofluorescence. Scale bar = 10 μm. **b.** U2OS stained with indicated SUMO1 MAb (red) and PML (green). Mouse and Rat SUMO1 MAbs (21C7, 4D12, 5B12, BD8B16, D11 and 852620) were co-stained with Rabbit PML antibody and Rabbit SUMO1 MAbs were co-stained with Mouse PML antibody. Scale bar = 10 μm. **c.** As for 5b but with SUMO2/3 MAbs **d.** As for 5b but with SUMO4 MAbs **e.** Quantification of SUMO1 MAb intensity signal per cell treated with either control (siNTC) or SUMO1 (siSUMO1) siRNA for 72 hours prior to fixation. Two tailed t-test denotes statistical differences between siNTC and siSUMO1. N= ~100 cells. **f.** As for 5e but using SUMO2/3 MAbs and SUMO2+3 siRNA. **g.** As for 5e but using SUMO4 MAbs and SUMO2+3 and SUMO4 siRNA.

### SUMO MAbs as enrichment agents

A common methodology for detecting SUMO conjugation of individual substrates is to use a SUMO antibody as an enrichment tool followed by immunoblotting against the studied substrate. Given the variability in sensitivity and specificity described herein, we set out to determine the ability of the clones to purify SUMOylated substrates by immunoprecipitation (IP). Using FLAG-SUMO complemented cells, we were able to compare IP efficiency from multiple antibodies in parallel using anti-FLAG M2. We used a modified version of the denaturing IP protocol developed by Becker et al (Becker et al., 2013).

In the investigation of IP by SUMO1 antibodies, we noted clones BD8B16 and D11 immunoprecipitated FLAG-SUMO1 to levels equivalent to FLAG M2. Two SUMO1 MAbs that performed poorly in western blot analysis (5B12 and 852620) nevertheless immunoprecipitated FLAG-SUMO1 (Figure 6a & b). We assessed the ability to enrich SUMO1 modified RANGAP1 and found C9H1, D11, EP298 and Y299 precipitated SUMO1-RANGAP1 as well or better than FLAG M2, while 21C7, 4D12, 5B12 and 852620 were 50% as efficient (Figure 6a & c).

**Figure 6.**
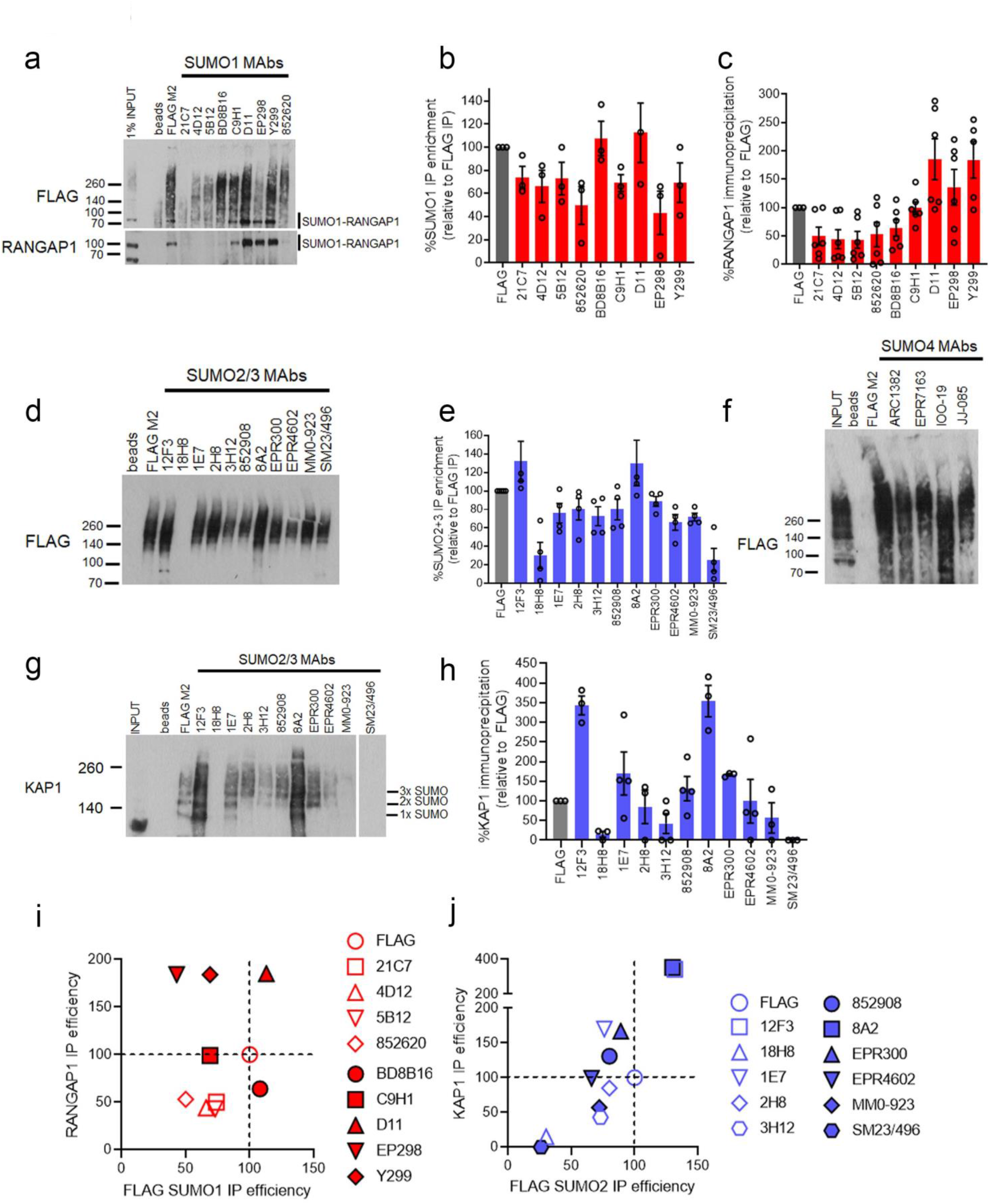
**a.** U2OS FLAG-SUMO1 treated with siSUMO1 and doxycycline for 48 hours prior to denaturing lysis and immunoprecipitation with indicated MAbs. Immunoprecipitated material was divided in two and probed with FLAG or RANGAP1 antibody. FLAG panel is from a 30 second exposure while RANGAP1 is 1 minute. The input for RANGAP1 detects both the unmodified and SUMO1 conjugated species (~70 and 100 kDa respectively). **b.** Quantitation of FLAG-SUMO1 IP signal (between 70 KDa and the well front) from 6 experiments. The enrichment is shown as a % relative to that of the M2 FLAG MAb as an internal control for each experiment. Error bars show SEM. **c.** Quantitation of RANGAP1 IP signal (100 kDa species) as for 6b. **d.** U2OS FLAG-SUMO2 treated as for 6a. **e.** Quantification of FLAG-SUMO2 enrichment as for 6b from 5 experiments. **f.** As for 6d but using SUMO4 MAbs. **g.** Immunoprecipitates from 6d were in parallel probed for KAP1. The approximate location of mono, di and tri-SUMO modified KAP1 is shown. **h.** Quantification of KAP1 enrichment by SUMO2/3 MAbs from 3 experiments as for 6c. **i.** Relative IP efficiencies for FLAG-SUMO1 (total SUMOylation) versus RANGAP1-SUMO1 enrichment. The IP efficiency of FLAG M2 is set at 100% for both conditions. **j.** As for 6i but with FLAG-SUMO2 and KAP1.

In the investigation of IP by SUMO2/3 antibodies, all but clones 18H8 and SM23/496 performed as well as M2 for SUMO conjugate immunoprecipitation (Figure 6d & e). All SUMO4 MAbs immunoprecipitated FLAG-SUMO2 (Figure 6f). We tested the ability of each to enrich for KAP1/TRIM28, a protein that has high basal autoSUMOylation in several cell lines and is SUMOylated as a monomer, multi-monomer, and polymer (Ivanov et al., 2007, Bruderer et al., 2011, Becker et al., 2013, Garvin et al., 2013) (Figure 6g & h). Clones 12F3 and 8A2 exceeded the ability of FLAG M2 to enrich SUMOylated KAP1. Mono-SUMOylated KAP1 was only detected by FLAG, 12F3, 1E7 and 8A2 (Figure 6g).

## Discussion

This study has explored the specificity and sensitivity of 24 antibodies raised to SUMO isoforms. We have tested them against monomeric and conjugated SUMO, in dot-blot, immunoblot, immunofluorescence and immunoprecipitation formats. We describe a tremendous variability in each antibody to detect SUMO states in these formats, with very few ‘all-rounders’ (Figure 7). Several companies have committed to improved antibody validation. Abcam, Cytoskeleton and BIORAD have validated their SUMO1 MAbs using *SUMO1* knock-out HAP1 cells. Our cataloguing of nine antibodies raised to SUMO1 finds that several are specific and sensitive in at least one detection format tested. SUMO1 MAb 852620 (R&D A-716) was entirely unspecific by immunoblot but was able to detect SUMO1 in dot blot and immunoprecipitation.. SUMO1 5B12 (MBL Life science) performed poorly in most applications except immunoprecipitation. Two SUMO1 MAbs (EP298 Abcam and D11 Santa Cruz) detect multiple nonspecific bands by immunoblot but performed well in other applications. The ability of SUMO1 MAbs to enrich SUMO1 modified RANGAP1 varied considerably. We also note that overall ability to enrich SUMO1 modified proteins does not necessarily correlate the ability to enrich RANGAP1 (Figure 7) presumably due to the accessibility of epitopes within modified proteins and complexes.

**Figure 7.**
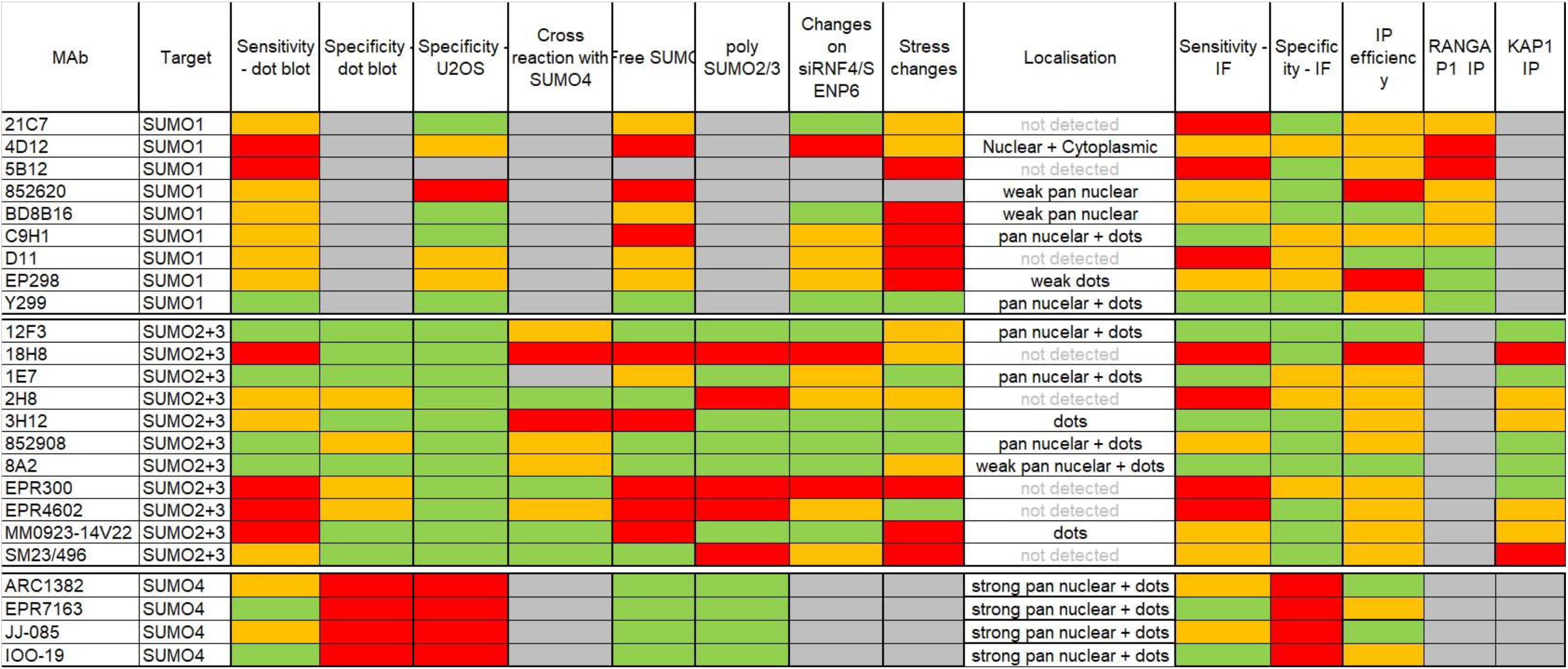
Summary of SUMO MAbs, grey boxes indicate condition not tested. The performance of each MAb for stated application was scored as high (green), moderate (orange), or low (red) using the following criteria. See Material and Methods, “Relative Quantification” for further definitions.

As SUMO2 is essential in most cell lines (McFarland et al., 2018) and mice (Wang et al., 2014), knockout cells are not commercially available, but have been recently described in U2OS (Bouchard et al., 2021). Our data suggest that many antibodies raised to SUMO2 and SUMO3 are sensitive in some of the detection formats we tested, although different antibodies are preferable for certain modalities (Figure 7). Many detect SUMO4 when it is overexpressed (Supplemental Figure 2c) but showed a preference for SUMO2/3 over SUMO4. We found that the ability to detect high molecular weight recombinant SUMO2 polymers did not necessarily correlate with the ability to detect changes in high molecular weight SUMO2/3 following RNF4 or SENP6 depletion (e.g., 1E7 and MM0-923 and SM23/496), presumably reflecting the greater complexity of SUMOylation in cells. It should be noted that like Ubiquitin, SUMO1-3 form polymers on multiple internal lysines with evidence of branching and mixed chains composed of other Ubls (Hendriks et al., 2018). An analysis of Ubiquitin antibodies highlight the variability of these reagents to detect different homotypic linkages (Emmerich and Cohen, 2015). It therefore seems likely that the same is true for SUMO MAbs. We also highlight the variation (0-15% of SUMO) in detection of monomeric / unconjugated SUMO by the tested MAbs (Figure 3a) and the inability of some clones (e.g. 4D12, EPR300, EPR4602) to detect monomeric SUMO by dot blot (Figure 1b-c).

Unlike RANGAP-1, KAP1 immunoprecipitation tracked with overall SUMO2/3 enrichment. Intriguingly at least two SUMO2/3 MAbs (12F3 and 8A2) outperformed FLAG M2 at immunoprecipitation. One potential reason for this is a partial knockdown of SUMO2/3 in our cells which would enable SUMO2/3 MAbs to detect both endogenous SUMO2/3 and exogenous FLAG-SUMO2 species, while the M2 MAb only enriches exogenous SUMO2.

All tested antibodies raised to SUMO4 failed to show specificity towards SUMO4 over the closely related SUMO2/3. In some formats, they showed greater selectivity for SUMO2/3 over SUMO4. Unlike other isoforms, SUMO4 is both nuclear and cytoplasmic (Figure 5a) and shows no increase in conjugation following cellular stresses (Figure 4). During the preparation of this report, clone EPR7163, raised against SUMO4, was rebranded as a SUMO2/3/4 antibody after in-house validation by Abcam. Our data suggests reports of specific SUMO4 biology using several of the antibodies raised to SUMO4 should be treated with caution. We tested 4 of 9 SUMO4 MAbs on the market. Those we did not test are unvalidated (2E4, 9H10-e), show nuclear staining not consistent with SUMO4 (3C3) or are classified as pan SUMO2/3/4 (C3).

Our catalogue of these essential tools reveals a subset that are robust and specific for the use of SUMO isoform detection in various methodologies.

## The authors declare no conflict of interest

### Author Contributions

AJG designed the study, undertook all blots, cell work, imaging and analysis, AL generated purified SUMO isoforms, JRM oversaw the study. AJG and JRM co-wrote the paper. All Authors have commented and edited the manuscript.

## Acknowledgements

Grant funding. Wellcome Trust 206343/Z/17/Z (A.J.G), University of Birmingham (A.L). We thank the Microscopy and Imaging Services at Birmingham University (MISBU) in the Tech Hub facility for microscope support and maintenance.

## Materials and Methods

### Recombinant SUMO proteins

Human SUMO1, SUMO2, and SUMO3 cDNA were cloned into pGEX4-T1 using BamHI and EcoR1 restriction sites and SUMO4 was cloned into pGEX6P1 using BamHI and EcoR1. Plasmids were transformed into BL21 DE3 (NEB) and clones were picked for growth in 40 mL LB (Melford) starter cultures supplemented with ampicillin (100 mg/ml) for overnight growth at 37 °C. For each litre of culture 10 mL of starter culture was added and grown at 37 °C/180 rpm to OD_595_ of 0.6. Temperature was reduced to 18 °C and 0.5 mM IPTG added for a 16-hour induction. Bacteria were pelleted at 5,000xg (4 °C) for 10 minutes and resuspended in lysis buffer (20 mM Tris-HCl pH 8, 130 mM NaCl, 1 mM EDTA, 1 mM EGTA, 1.5 mM MgCl_2_, 1% Triton X-100, 10% Glycerol, 1 mM DTT, Roche complete protease inhibitor) with 0.5 mg/mL Lysozyme and placed on a roller for 45 minutes at 4 °C. Lysates were sonicated five times for 30s followed by centrifugation (48,000xg at 4 °C for 30 minutes) and passing supernatant through a 0.45 μm syringe filter. Lysates were incubated with 250 μL of GST Sepharose 4B (GE Healthcare) for 3 hours at 4 °C. Beads were pelleted and washed three times in 10 mL lysis buffer and once in 10 mL cleavage buffer. For SUMO1-3 thrombin (16 U; Promega) was used for cleavage in buffer (20 mM Tris-HCl pH 8.4, 150 mM NaCl, 1.5 mM CaCl_2_), while PreScission Protease (30 U; SIGMA) was used for SUMO4 owing to a partial thrombin cleavage site unique to SUMO4, buffer (50 mM Tris-HCl pH 7, 150 mM NaCl, 1 mM EDTA, 1mM DTT). 500 μL buffer was added to GST-SUMO1-4 beads with respective cleavage protease and incubated for 16 hours at 4 °C with agitation. GST beads were pelleted with centrifugation at 1,000xg at 4 °C for 3 minutes and supernatant collected for further centrifugation at 14,000xg (4 °C) for 20 minutes. Samples were passed through a Superdex Increase 10/300 GL column attached to an AKTA pure (UNICORN software) for size-exclusion chromatography equilibrated in 20 mM Hepes pH7.5, 100 mM NaCl, 0.5 mM TCEP. SUMO protein fractions were pooled and aliquoted for storage at −80 °C. SUMO2 dimers (ULC-200-0505) and 2-8 polymers (ULC-210-025) were purchased from R&D.

### Western blots

Unless otherwise stated all cell lysates were made by direct lysis in 6X Laemmli buffer (Alfa Aesar) to inactive SENPs and denature proteins followed by heating at 95 °C for 5 minutes, sonication and centrifugation. Subcellular fractions were generated according to (Garvin et al., 2019a). Lysates were separated on 4-20% Tris-Glycine Novex wedgewell gels (Invitrogen) followed by wet transfer (stacking and resolving gel) in transfer supplemented with 20% methanol on 0.45mm PVDF membrane (IMMOBILON-P, Millipore). PVDF was selected over nitrocellulose as it shows an improved transfer of free SUMO (Castoralova et al., 2012). PVDF membranes were stained with Ponceau S stain (Merck) to confirm full transfer followed by blocking for 30 minutes at room temperature in 5% non-fat milk in PBST (0.01% Triton-X100). Membranes were probed with primary antibody (1 mg/mL) with gentle rolling at 4 °C for 16 hours followed by 3x 10 minutes washes with PBST and incubation with secondary HRP antibodies (1:10000) in 5% milk/PBST for 2 hours at room temperature. Following 3x 10-minute PBST washes membranes were exposed to ECL reagent (EZ-ECL, GeneFlow) for two minutes prior to exposure with blue X-ray film (Wolflabs).

### Dot blots

Recombinant SUMO (rSUMO) proteins were diluted to 3 μg/mL in 6X Laemmli buffer and diluted serially 1:2 10 times to a final dilution of 3 ng/ mL. After methanol activation, PVDF membranes were dried and dotted with 1 mL of denatured rSUMO. After drying at room temperature for 30 minutes blots were probed as for western blots. Each blot was exposed to X-Ray film at four exposure time points (0.5, 1, 2 and 10 minutes).

### Peptide competition

For peptide competition assays 12F3, 2H8, 852908, 8A2, 21C7, ARC1382, EPR7163, JJ-085, IOO-19, 5B12, BD8B16, C9H1, D11, Y299 and MM0-923 were measured by peptide competition dot blot as previously described (Becker et al., 2013). As some MAbs performed poorly by dot blot (EPR4602, EPR300) the same protocol was adapted to western blot competitions with U2OS lysates separated on 6% SDS-PAGE gels using the same concentration of peptide as for dot blots.

### Cell culture

Culture conditions for cell lines are shown in Supplemental Table 5. 6XHis-FLAG SUMO1, SUMO2 and HA-FLAG SUMO4 FlpIn stable cell lines were generated using U2OS^TrEx-FlpIn^ (a gift from Grant Stewart, University of Birmingham) cells transfected with pcDNA5/FRT/TO based vectors (Supplemental table 2) and the recombinase pOG44 (Invitrogen) using FuGene6 (Promega) at a ratio of 4 ml FuGENE/ mg DNA. After 48 hr, cells were placed into hygromycin selection media (100 μg/ml) and grown until colonies formed on plasmid-transfected plates but not controls.

### Immunofluorescent

Cells were plated at 2.5 x 10^4^ cells/well on 13 mm glass coverslips in 24 well plates (Corning) and attached overnight prior to siRNA depletion for 48 hours. For pre-extraction after 1x PBS wash cells were treated with 250 mL / well ice cold CSK buffer (100 mM NaCl, 300 mM sucrose, 3 mM MgCl2, 0.7% Triton-X100 and 10 mM PIPES) for 30 seconds prior to fixation with 4% Paraformaldehyde (PFA) in PBS at room temperature for 10 minutes. For non-pre-extracted samples cells were fixed in 4% PFA at room temperature for 10 minutes followed by permeabilisation with 0.5% Triton in PBS for 5 minutes. Coverslips were blocked with 5% FBS in PBS for 1 hour at room temperature followed by incubation with primary antibodies at 1 mg/mL (or 1:1000 for recombinant and CST MAbs) overnight at 4°C in 5% FBS. Coverslips were washed twice with PBS followed by incubation with Alexa-Fluor 555 conjugated secondary antibodies at 1:2500 for 2 hours at room temperature in the dark. Cells were washed twice with PBS prior to incubation with 250 μL of Hoechst (1 μg/mL) for 2 minutes. Coverslips were mounted on slides using Immuno-Mount (Thermo Scientific) and sealed. Imaging was carried out on a Leica DM6000B microscope using an HBO lamp with 100W mercury short arc UV bulb light source. Images were captured at each wavelength sequentially using the Plan Apochromat HCX 100x/1.4 Oil objective at a resolution of 1392×1040 pixels.

### Denaturing SUMO Immunoprecipitations

U2OS FlpIn 6xHIS-FLAG SUMO1 or SUMO2 cells were plated on 15 cm^2^ plates at 5×10^6^ for 24 hours prior to siRNA depletion with either 10 nM SUMO1 or 10 nM each of SUMO2 and SUMO3 siRNA (SIGMA). Doxycycline (1 mg/mL) was added concurrently with siRNA depletion for 48 hours. Cells were trypsinised and pelleted in cold PBS followed by lysis in 1% SDS Lysis buffer (20 mM Sodium phosphate pH 7.4, 150 mM NaCl, 1% SDS, 1% Triton, 0.5% Sodium deoxycholate, 5 mM EDTA, 200 mM IAA and complete Protease and Phosphatase inhibitors). Lysates were sonicated 3x for 20 seconds followed by the addition of 50 mM DTT and boiling for 10 minutes. Lysates were clarified by centrifugation and frozen at −80 °C for later use. For immunoprecipitation, Protein A/G Plus agarose beads (Pierce) were washed twice in 10x packed bead volumes of RIPA buffer (20 mM Sodium phosphate pH 7.4, 150 mM NaCl, 1% Triton, 0.5% Sodium deoxycholate, 5 mM EDTA, 100 mM NEM and Protease / Phosphatase inhibitor cocktail). Antibodies were bound at 4 mg / 30 μL of beads at 4 °C for 3 hours. 1% SDS lysates were thawed and diluted 10x with RIPA buffer before incubating 1 mL of lysate with MAb conjugated beads overnight at 4 °C. A fraction of lysate was retained as input, and the beads were washed 3x with 1 mL of RIPA buffer before eluting in 30 μL of 6X Laemmli buffer and boiling.

### Statistics

Unless otherwise stated all statistical analysis was by two-sided Students T-test throughout. *<p0.05, **p<0.01, ***P<0.005 ****P<0.001. All centre values are given as the mean and all error bars are standard error about the mean (s.e.m). Data was analysed using GraphPad Prism 7.03.

### Quantification

All Western Blot or Image analysis for quantification was done using ImageJ unless otherwise specified.

### Relative quantification (Figure 7)

**Sensitivity by dot blot**, MAbs that produced a saturating signal (less than 25% change between two exposure time points) across at least the five most concentrated dots at the 10-minute exposure were classed as having high sensitivity by dot blot. Those that produced a saturating signal on at least two concentrations were classed as moderate. MAbs that did not reach saturation at any concentration or exposure time were classed as low.

**Specificity by dot blot**, SUMO2/3 or SUMO4 MAbs that did not produce any signal for the reciprocal isoform were classed as highly specific by dot blot. Those that produce any signal were moderate and those that detected signal at the 0.5-minute exposure on the reciprocal isoform blot were classed as low specificity.

**Specificity in U2OS** was classed as high if the signal from either the siNTC or complemented lanes were at least 4x higher than the siSUMO lane. Moderate specificity was classed as any nonspecific bands detected by the MAb on siSUMO treatment. Low specificity was for MAbs that showed no difference between control and siSUMO treatment.

**Cross reaction with SUMO4 in U2OS** was assessed by detection of HA-FLAG SUMO4 overexpression. Where there was less than a 20% difference between FLAG and SUMO MAb detection of SUMO4 the MAb was classed as highly cross reactive, moderately cross reactive if the MAb detected 50% or less of HA-FLAG SUMO4 and low cross reactive if less than 20%. Detection of free SUMO was measured relative to the FLAG-SUMO, levels of more than 80% of FLAG were considered high, more than 50% moderate and less than 20% low.

**PolySUMO2 detection** was measured as the ability to detect SUMO2 species higher than a tetramer at 2 minutes exposure. MAbs were either scored as high (tetramers and above detected) or low (not detectable).

**Changes in SUMO conjugates on RNF4 or SENP6 siRNA depletion** were assessed as high if either RNF4 or SENP6 depletion caused a >100% increase in conjugate levels versus FLAG-SUMO, moderate for >50% increase or low for less than 10%.

**Stress changes** were assessed across all conditions versus FLAG SUMO1 or FLAG SUMO2. Any change in stress conjugates that average (across all stresses) as equal to or greater than the FLAG-SUMO control was considered sensitive, those within 20% of FLAG were considered moderately sensitive to stress and those less than 20% were classed as low sensitivity.

**Sensitivity by immunofluorescence** was measured on the mean fluorescence intensity in siNTC cells (Arbitrary units). Less than 2000 was classed as low sensitivity, between 2000 and 4000 was classed as moderate sensitivity and higher than 4000 was classed as high sensitivity.

**Specificity by immunofluorescence** was classed as high if signal between siNTC and siSUMO was statistically significant (two tailed students t-test) in both pre-extracted and non-pre-extracted conditions, moderate if significance was only detected in one of the extraction conditions and low if signal was not significantly reduced under either condition.

**IP efficiency** was measured relative to the FLAG-SUMO isoform control. High efficiency was classed if 100% or more of the IP signal was detected, moderate if 50% and low if less than 25%.

**RANGAP1 and KAP1 IP efficiency** was measured as for total SUMO IP efficiency.

---

**Supplemental Figure 1.**
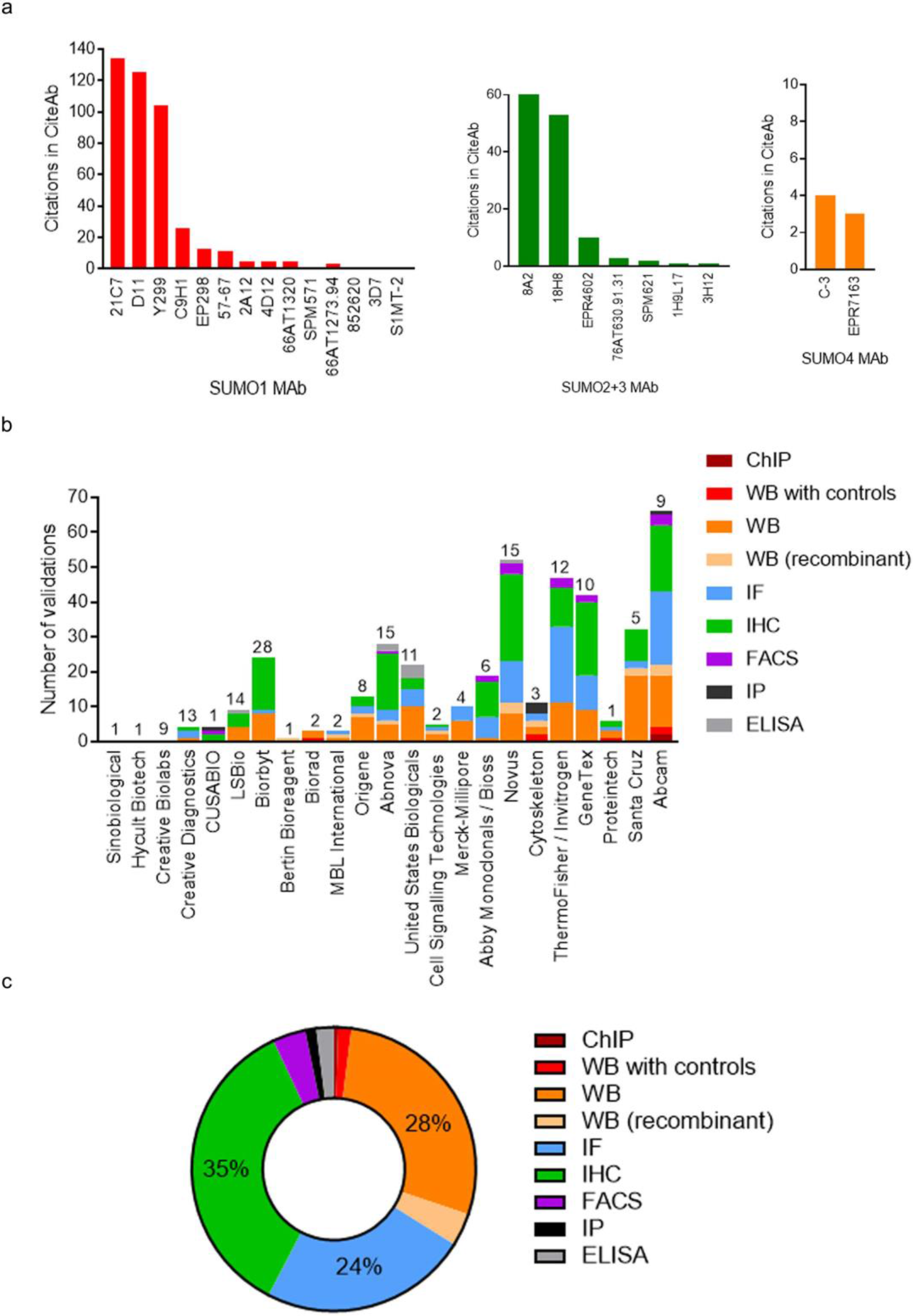
**a.** Citations for each SUMO MAb from the CiteAb database as of March 2022. **b.** Total number of validations for SUMO1, SUMO2/3 and SUMO4 MAbs ranked by vendor. The total number of SUMO MAbs available per vendor (as of March 2022) is shown above each bar. Validations include any supporting images or data available on the products webpage including those generated by the vendor. Reviews on the product webpage were only included if they contained accompanying images or data. Types of validation are colour coded. **c.** Type of validation across all SUMO MAbs and vendors. Controls include either negative (knockdown or knockout) or positive (over-expression or induced expression with stress).

**Supplemental Figure 2.**
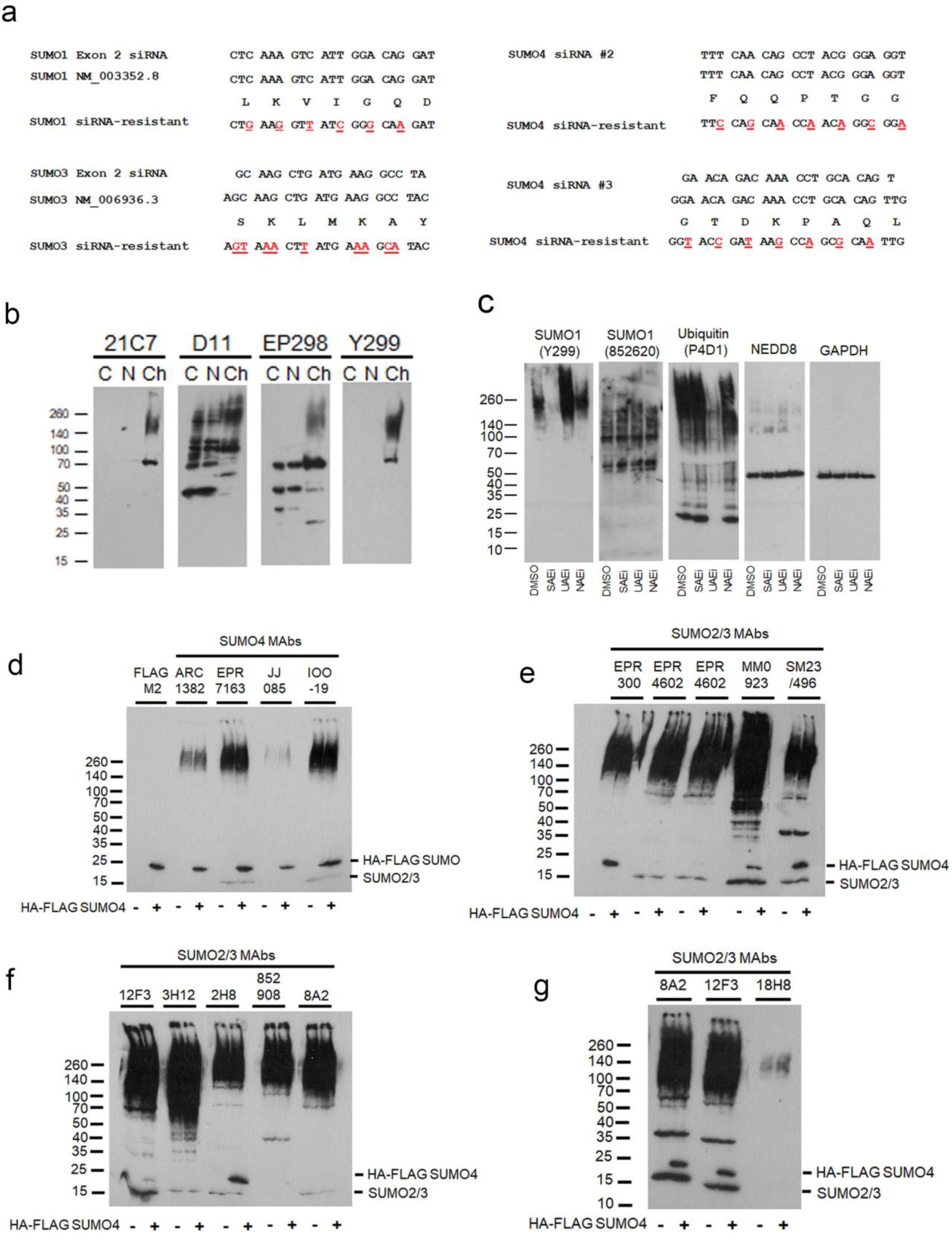
**a.** Location of synonymous mutations in SUMO1, SUMO3 and SUMO4 cDNA to generate siRNA resistant species. **b.** U2OS lysates separated by fractionation into cytoplasm (C), soluble nuclear (N) or chromatin (Ch) were probed with 21C7 and Y299 control SUMO1 MAbs and two SUMO1 MAbs that detect non-specific bands (D11 and EP298). ChIP (Chromatin Immunoprecipitation), WB (Western blot), recombinant – testing on E. coli derived SUMO proteins, IF (Immunofluorescence), IHC (Immunohistochemistry), FACS (Fluorescence Activated Cell Sorting), IP (Immunoprecipitation), ELISA (enzyme-linked immunosorbent assay). **c.** U2OS treated for six hours with 10 μM each of ML-792 (SUMO E1 inhibitor), TAK-243 (Ubiquitin E1 inhibitor) or MLN4924 (NEDD8 E1 inhibitor) prior to lysis (400 μL Laemmli buffer / 6 well), separation on 4-20% SDS PAGE gels (20 μL / lane). **d, e, and f**. U2OS cells expressing HA-FLAG SUMO4 untreated (−) or dox treated (+) for 48 hours prior to lysis in Laemmli buffer. PVDF membranes divided and probed with indicated MAbs but imaged side by side for comparison. Two-minute film exposures shown. **g**. As for 2d-f but 10-minute exposure shown.

**Supplemental Figure 3a.**
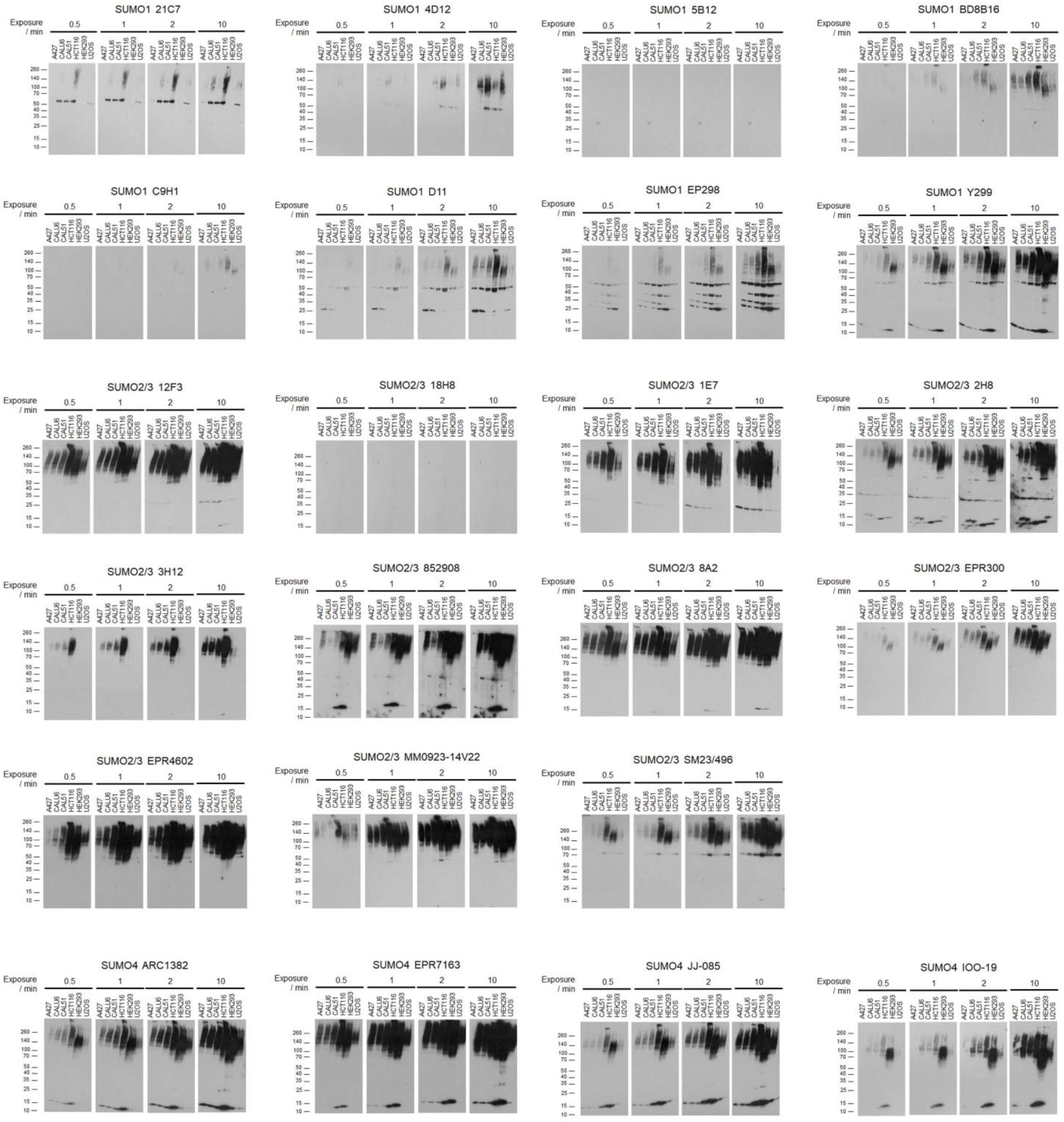
All exposures of immunoblots related to Figure 3a and 3b

**Supplemental Figure 3b.**
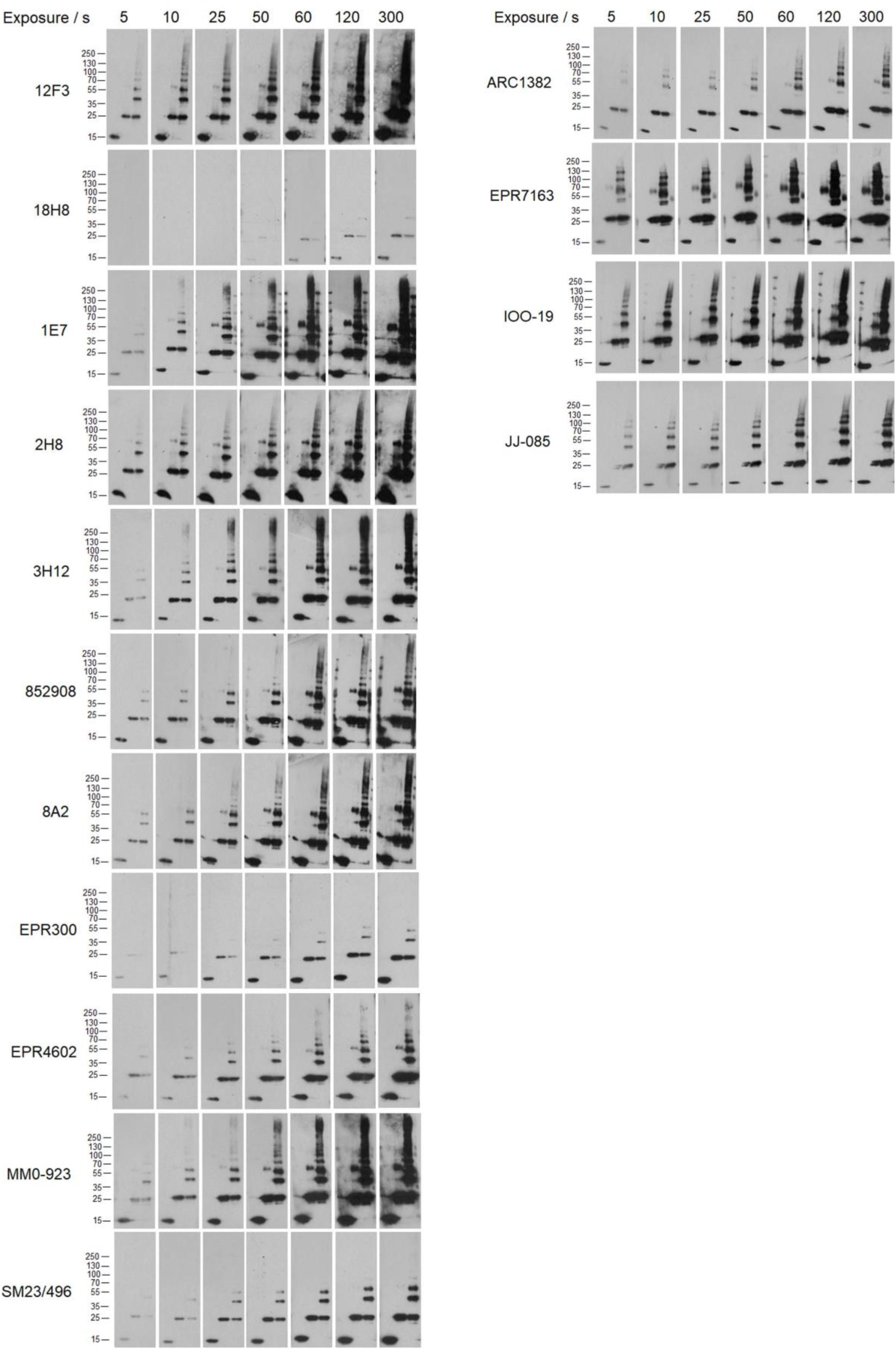
All exposures of immunoblots related to Figure 3c

**Supplemental Figure 4a.**
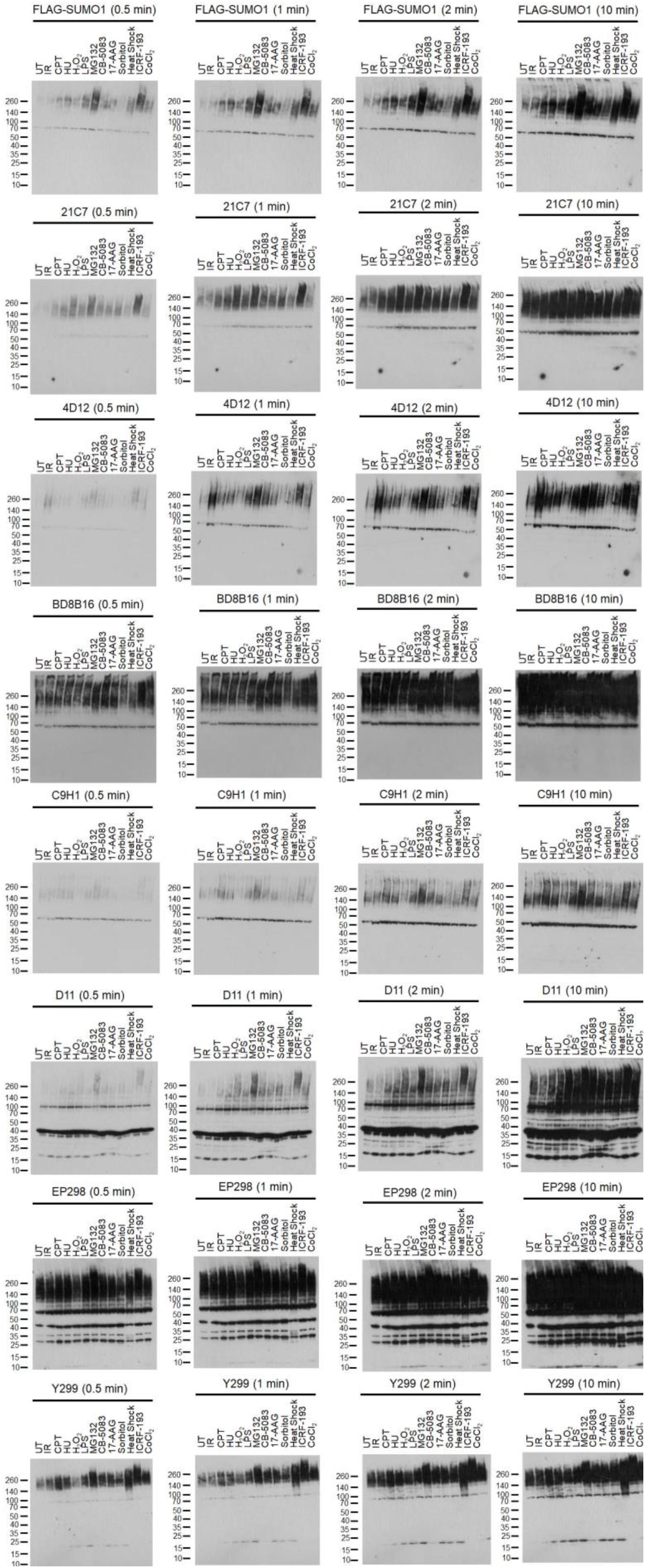
All exposures of SUMO1 immunoblots related to figure 4b.

**Supplemental Figure 4b.**
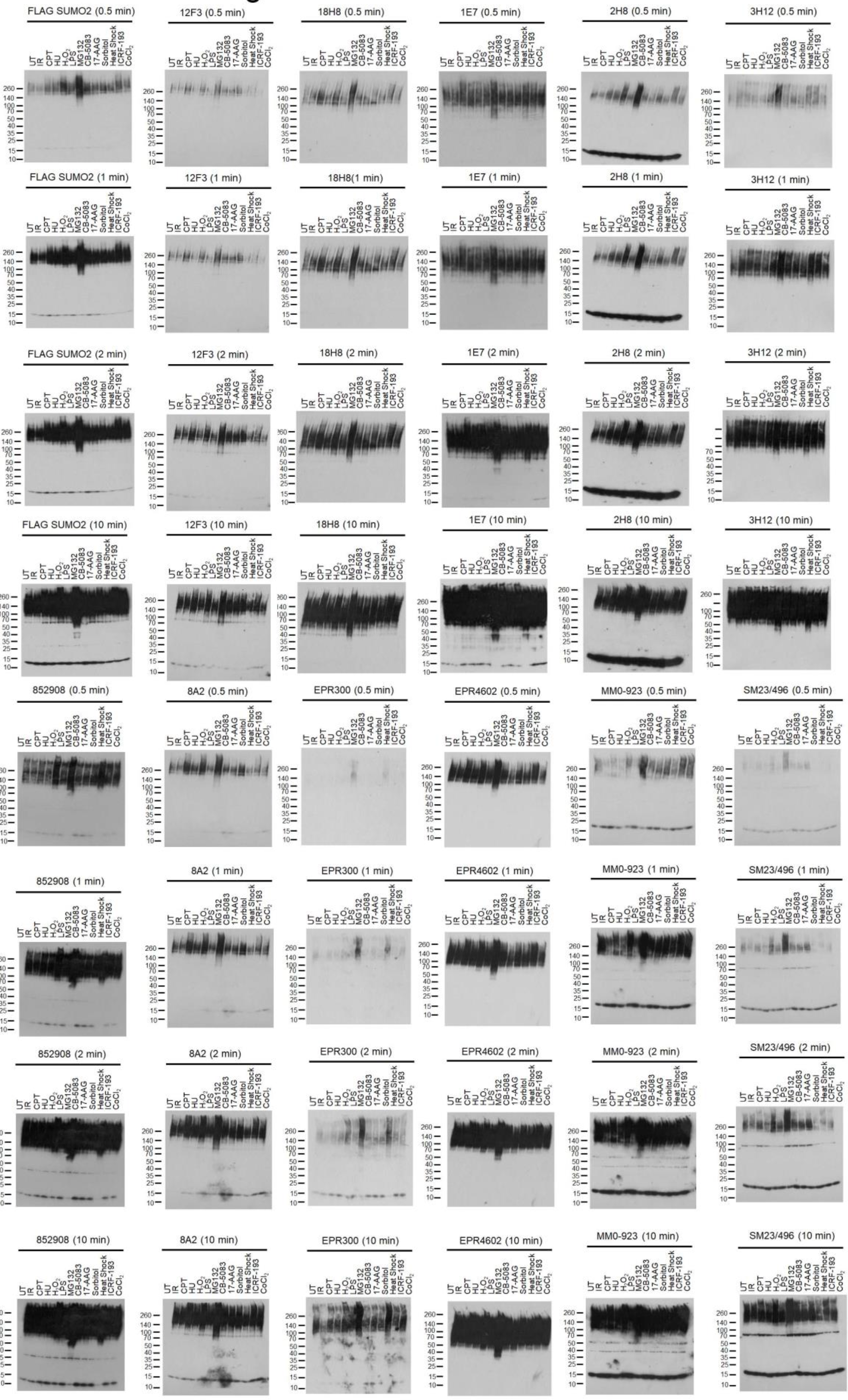
All exposures of SUMO2/3 immunoblot related to Figure 4b.

**Supplemental Figure 4c.**
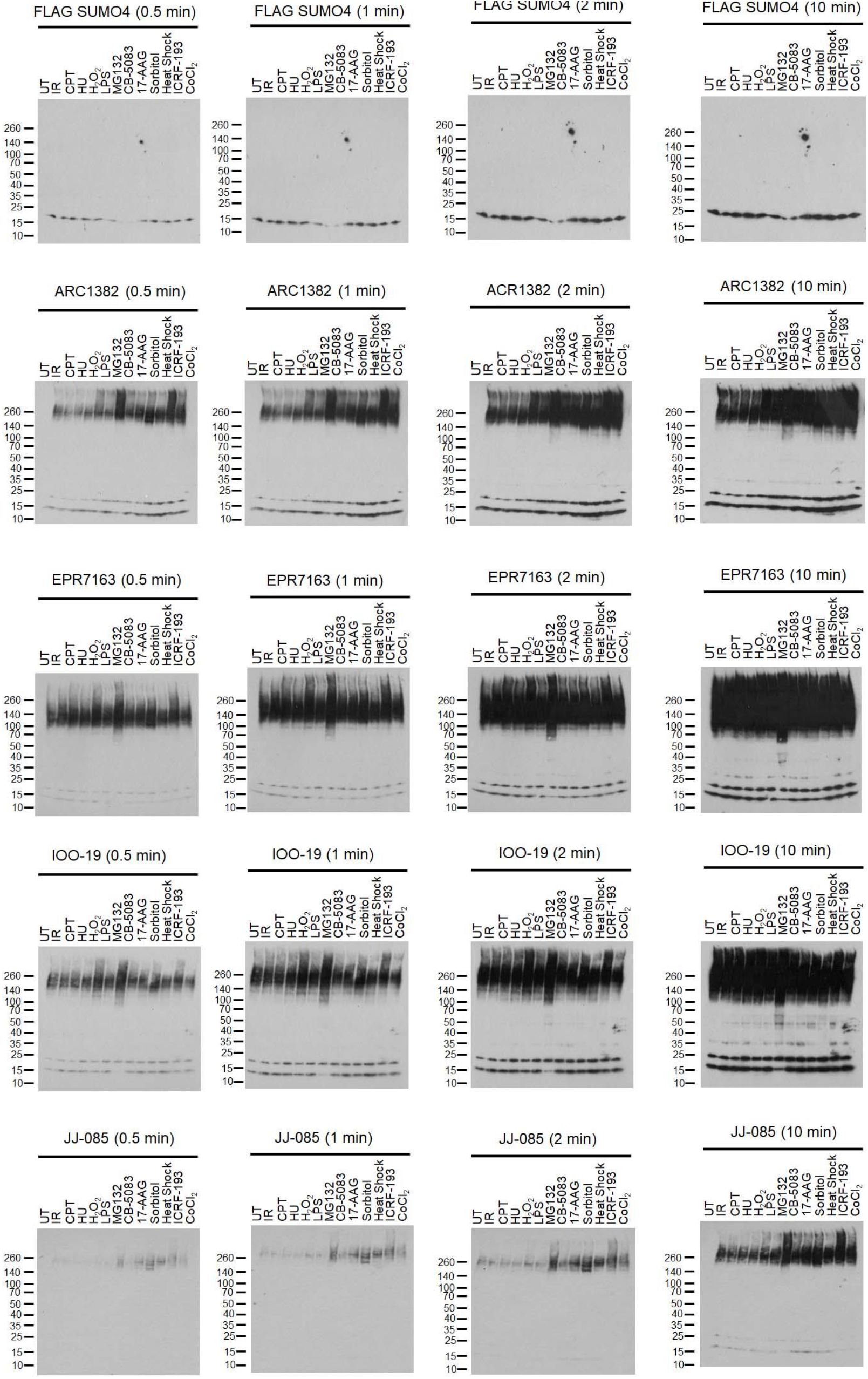
All exposures of SUMO4 immunoblot related to Figure 4d.

**Supplemental Figure 5.**
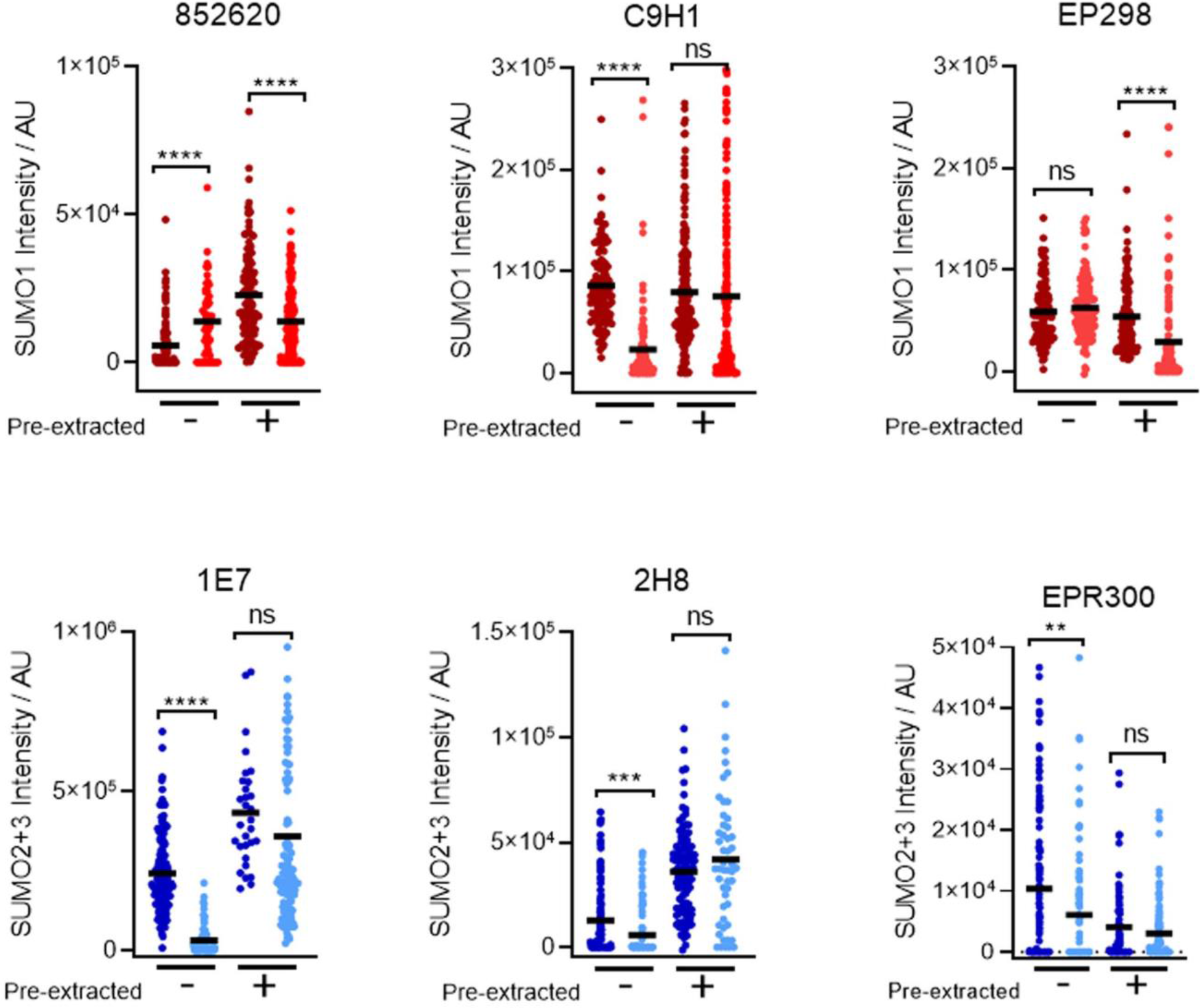
**a**. Immunofluorescent intensity (AU) of indicated SUMO MAb in U2OS cells fixed with PFA prior to Triton X-100 permeabilisation (−) or pre-extracted with CSK buffer prior to fixation (+). Statistical significance by two tailed students t-test is indicated. Ns = not significant. N= ~100 cells.

## Supplemental Tables

**Supplemental Table 2.**

| **Plasmid Name** | **Source** |
| --- | --- |
| 6XHis-FLAG SUMO1 WT | This paper / GenScript |
| 6XHis-FLAG SUMO2 WT | This paper / GenScript |
| 6XHis-FLAG SUMO3 WT | This paper / GenScript |
| HA-FLAG SUMO4 WT | This paper / GenScript |
| HA-FLAG SUMO4 M55V | This paper / GenScript |
| pGEX-4T1 ProSUMO1 | This paper / GenScript |
| pGEX-4T1 SUMO1 WT | This paper / GenScript |
| pGEX-4T1 ProSUMO2 | This paper / GenScript |
| pGEX-4T1 SUMO2 WT | This paper / GenScript |
| pGEX-4T1 ProSUMO3 | This paper / GenScript |
| pGEX-4T1 SUMO3 WT | This paper / GenScript |
| pGEX-6P1 SUMO4 WT | This paper / GenScript |
| pGEX-6P1 SUMO4 M55V | This paper / GenScript |

**Supplemental Table 3.**

| **Antibody** | **Catalogue number** | **Supplier** | **Lot number** | **Concentration / Use** |
| --- | --- | --- | --- | --- |
| FLAG (Gt) | ab1257 | Abcam | GR225879-35 | 1:1000 (WB) |
| FLAG M2 (Ms) | F1804 | Sigma | SLCC6485 | 1:2000 (WB, IF) |
| GAPDH 6C5 (Ms) | CB1001 | Calbiochem | 3725987 | 1:5000 (WB) |
| HA.11 (Ms) | 901501 | Biolegend | B272772 | 1:1000 (WB, IF) |
| KAP1 (Ms) | NB100-47549 | Novus | VK307456 | 1:1000 (WB) |
| NEDD8 (Ms) | sc-373741 | Santa Cruz | J2816 | 1:200 (WB) |
| PML C7 (Ms) | ab96051 | Abcam | GR3284434-1 | 1:1000 (IF) |
| PML (Rb) | PA5-79835 | Invitrogen | VL31407955 | 1:1000 (IF) |
| RANGAP1 (Rb) | 36067S | CST | LOT1 | 1:1000 (WB) |
| RNF4 (Goat) | AF7964 | R&D | CIRA0217091 | 1:250 (WB) |
| SENP6 (Rb) | HPA024376 | Merck | A106728 | 1:1000 (WB) |
| SUMO1 Y299 (Rb) | ab32058 | Abcam | GR3244068-3 | 1:1000 (WB, IF) |
| SUMO1 4D12 (Rat) | MABS1223 | Millipore | 3540377 | 1:500 (WB, IF) |
| SUMO1 EP298 (Rb) | ab133352 | Abcam | GR268526-10 | 1:1000 (WB, IF) |
| SUMO1 5B12 (Ms) | M113-3 | MBL |  | 1:1000 (WB, IF) |
| SUMO1 D11 (Ms) | sc-9060 | Santa Cruz | A2021 | 1:200 (WB, IF) |
| SUMO1 852620 (Rat) | A-716 | R&D Systems |  | 1:500 (WB, IF) |
| SUMO1 5D8B16 (Ms) | ASM01 | Cytoskeleton | LOT 3 | 1:1000 (WB, IF) |
| SUMO1 C9H1 (Rb) | 4940S | CST | LOT 7 | 1:1000 (WB, IF) |
| SUMO1 21C7 (Ms) | 33-2411 | Invitrogen | WE326808 | 1:500 (WB, IF) |
| SUMO2/3 8A2 (Ms) | ab81371 | Abcam | GR3233360-1 | 1:1000 (WB, IF) |
| SUMO2/3 EPR4602 (Rb) | ab109005 | Abcam | GR154070-1 | 1:1000 (WB, IF) |
| SUMO2/3 3H12 (Rat) | ab191492 | Abcam |  | 1:1000 (WB, IF) |
| SUMO2/3 SM23/496 (Ms) | ab190757 | Abcam |  | 1:200 (WB, IF) |
| SUMO2/3 852908 (Rat) | A-718 | R&D Systems |  | 1:500 (WB, IF) |
| SUMO2/3 2H8 (Ms) | SAB1402359 | Sigma | 11238-2H8 | 1:500 (WB, IF) |
| SUMO2/3 12F3 (Ms) | ASM23 | Cytoskeleton | LOT 11 | 1:1000 (WB, IF) |
| SUMO2/3 18H8 (Rb) | 4971S | CST | LOT 7 | 1:1000 (WB, IF) |
| SUMO2/3 IE7 (Ms) | M114-3 | MBL | LOT 28 | 1:1000 (WB, IF) |
| SUMO2/3 EPR300 (Rb) | ab109196 | Abcam | GR96828-9 | 1:1000 (WB, IF) |
| SUMO2/3 MM0923-14V22 (Rat) | ab201594 | Abcam |  | 1:200 (WB, IF) |
| SUMO4 EPR7163 (Rb) | ab126606 | Abcam |  | 1:1000 (WB, IF) |
| SUMO4 JJ085-01 (Rb) | MA5-32478 | Invitrogen | WC32 16805 | 1:1000 (WB, IF) |
| SUMO4 ARC1382 (Rb) | MA535745 | Life Technologies | WC3224343A | 1:1000 (WB, IF) |
| SUMO4 IOO-19 (Rb) | M06740 | Boster Biological Technologies | BS21031540 | 1:1000 (WB, IF) |
| Ubiquitin P4D1 (Ms) | sc-8017 | Santa Cruz |  | 1:200 (WB) |
| Donkey anti-mouse-488 AF | A21202 | Invitrogen |  | 1:2000 (IF) |
| Donkey anti-rabbit-488 AF | A21206 | Invitrogen |  | 1:2000 (IF) |
| Donkey anti-mouse-555 AF | A31570 | Invitrogen |  | 1:2000 (IF) |
| Donkey anti-rabbit-555 AF | A31572 | Invitrogen |  | 1:2000 (IF) |
| Polyclonal Swine Anti-Rabbit Immunoglobulins/HRP | P021702-2 | Agilent Technologies |  | 1:5000 (WB) |
| Goat anti Rat HRP | AP136P | Merck |  | 1:5000 (WB) |
| Polyclonal Rabbit Anti-Mouse Immunoglobulins/HRP | P0161 | Agilent Technologies |  | 1:5000 (WB) |

**Supplemental Table 4.**
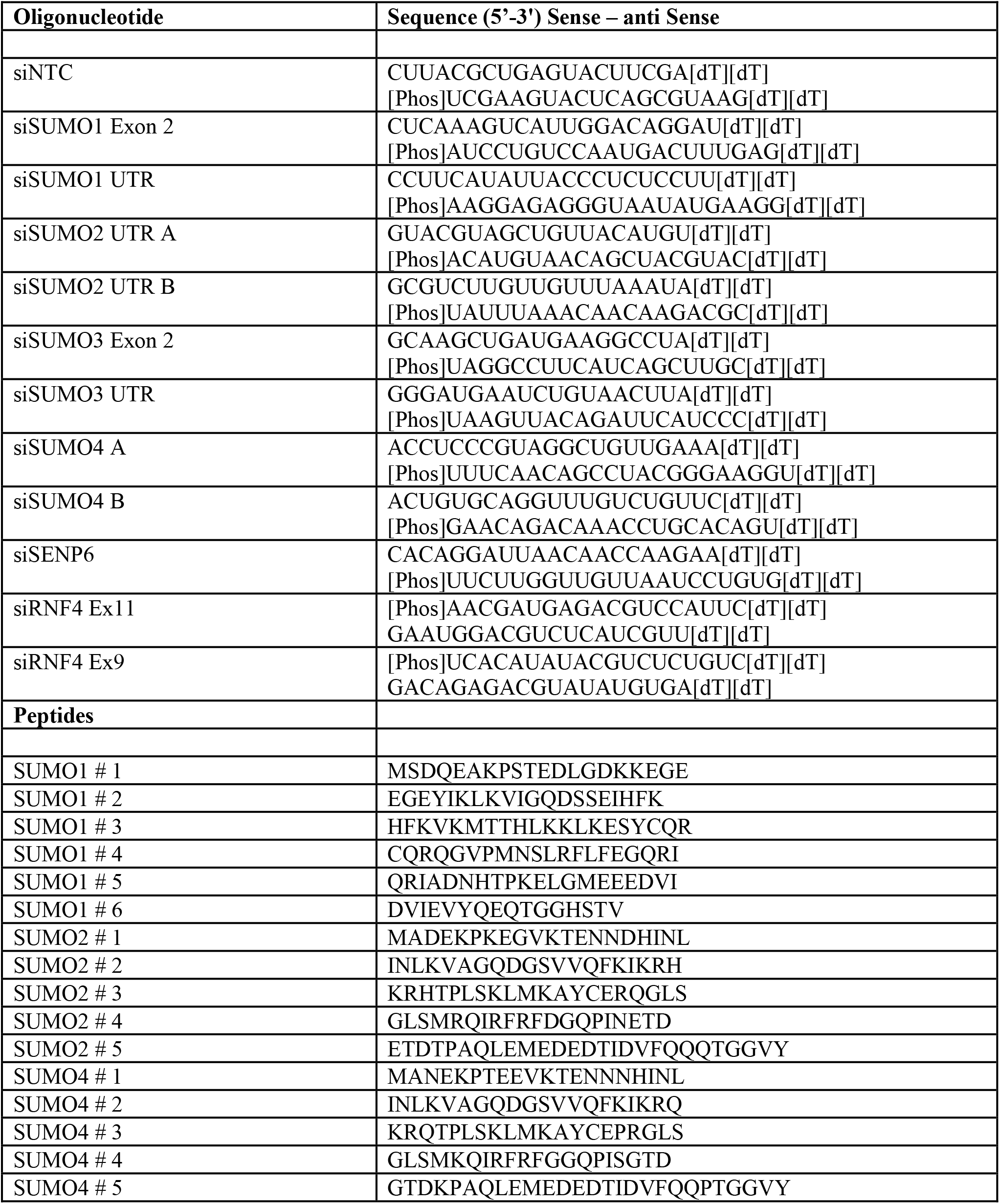

**Supplemental Table 5.**

| Cell line | Growth conditions | RRID |
| --- | --- | --- |
| U2OS FlpIn | DMEM, 10% FBS, 1% Penicillin/Streptomycin | N/A |
| U2OS FlpIn 6xHis-FLAG SUMO1 WT | DMEM, 10% FBS, 1% Penicillin/Streptomycin, 100 mg / mL Hygromycin B | N/A |
| U2OS FlpIn 6xHis-FLAG SUMO2 WT | DMEM, 10% FBS, 1% Penicillin/Streptomycin, 100 mg / mL Hygromycin B | N/A |
| U2OS FlpIn 6xHis-FLAG SUMO3 WT | DMEM, 10% FBS, 1% Penicillin/Streptomycin, 100 mg / mL Hygromycin B | N/A |
| U2OS FlpIn HA-FLAG SUMO4 WT | DMEM, 10% FBS, 1% Penicillin/Streptomycin, 100 mg / mL Hygromycin B | N/A |
| A427 | EMEM, 10% FBS, 1% Penicillin/Streptomycin, 1% NEAA, 1mM Sodium Pyruvate | CVCL_1055 |
| CALU6 | EMEM, 10% FBS, 1% Penicillin/Streptomycin, 1% NEAA, 1mM Sodium Pyruvate | CVCL_0236 |
| CAL51 | DMEM, 10% FBS, 1% Penicillin/Streptomycin | CVCL_1110 |
| HCT116 | DMEM, 10% FBS, 1% Penicillin/Streptomycin | CVCL_0291 |
| HEK293 TREx FlpIn | DMEM, 10% FBS, 1% Penicillin/Streptomycin | CVCL_U427 |

**Supplemental Table 6.**
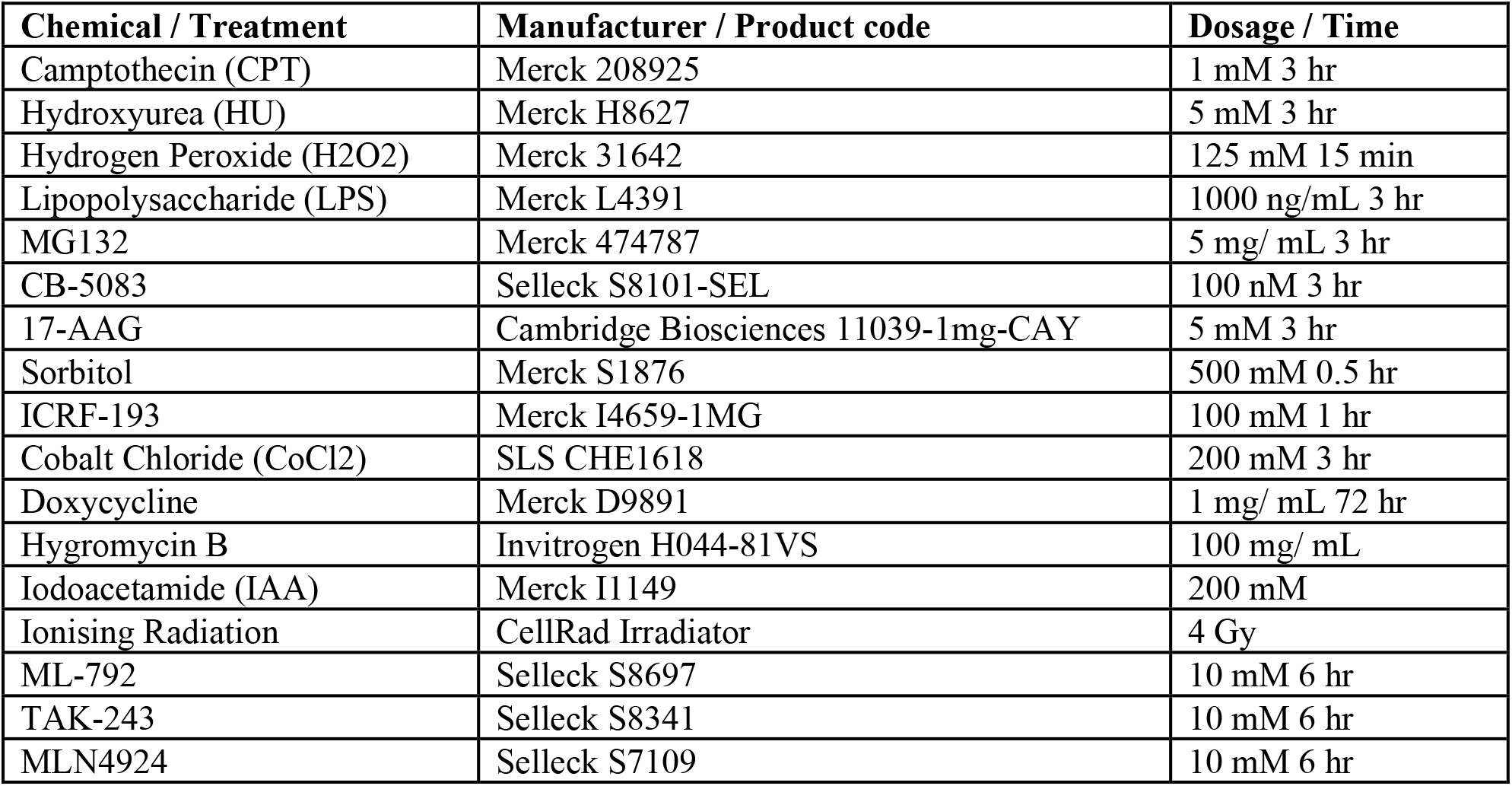

